# Prioritizing prognostic-associated subpopulations and individualized recurrence risk signatures from single-cell transcriptomes of colorectal cancer

**DOI:** 10.1101/2022.10.12.511912

**Authors:** Mengsha Tong, Yuxiang Lin, Wenxian Yang, Jinsheng Song, Zheyang Zhang, Jiajing Xie, Jingyi Tian, Shijie Luo, Chenyu Liang, Jialiang Huang, Rongshan Yu

**Affiliations:** State Key Laboratory of Cellular Stress Biology, School of Life Sciences, Faculty of Medicine and Life Sciences, Xiamen University, Xiamen, Fujian 361102, China; National Institute for Data Science in Health and Medicine, Xiamen University, Xiamen, Fujian 361102, China; School of Informatics, Xiamen University, Xiamen 316000, China; Aginome Scientific, Xiamen, Fujian 316005, China

**Keywords:** single-cell gene pair signatures, relative expression orderings, Colorectal cancer, recurrence risk signatures

## Abstract

Colorectal cancer (CRC) is one of the most common gastrointestinal malignancies. There are few recurrence risk signatures for CRC patients. Single-cell RNA-sequencing (scRNA-seq) provides a high resolution platform for prognostic signature detection. However, scRNA-seq is not practical in large cohorts due to its high cost and most single-cell experiments lack clinical phenotype information. Few studies have been reported to use external bulk transcriptome with survival time to guide the detection of key cell subtypes in scRNA-seq data. We proposed a data analysis framework to prioritize prognostic-associated subpopulations based on relative expression orderings (REOs). Cell type specific gene pairs (C-GPs) were identified to evaluate prognostic value for each cell type. We found REOs-based signatures could accurately classify most cell subtypes. C-GPs achieves higher precision compared with four current methods. Moreover, we developed single-cell gene pair signatures to predict recurrence risk for patients individually. Fibro_SGK1 cells and IgA+ IGLC2+ B cells were novel prognostic-associated subpopulations. A user-friendly toolkit, scRank^XMBD^ (https://github.com/xmuyulab/scRank-XMBD), was developed to enable implementation of this framework. Our work facilitate the application of the rank-based method in scRNA-seq data for prognostic biomarker discovery and precision oncology.

## Introduction

Colorectal cancer (CRC) is the third most common cancer and a leading cause of morbidity and mortality worldwide^1^. The implementation of curative resection and adjuvant chemotherapy has improved the overall prognosis of CRCs. However, about 25–40% of patients would relapse after primary radical resection^2^. The tumor-node-metastasis (TNM) staging system from International American Joint Committee on Cancer/Union for International Cancer Control (AJCC/UICC) remains the most important guideline for classifying patients and making therapeutic decisions. Unfortunately, due to high heterogeneity in CRC, the clinical outcomes for patients of the same stage can be very different^2^. Therefore, there is an urgent clinical need for molecular biomarkers that predict early relapse in CRC patients for more precise patient stratification.

The inter-patient heterogeneity of CRC has been revealed by genomic and epigenetic analysis, gene expression profiles and tumor microenvironment (TME). At the genetic level, several DNA biomarkers including microsatellite instability (MSI), BRAF and KRAS mutations, CpG island methylator phenotype (CIMP) and chromosomal instability (CIN), have been reported^3^. At the transcriptome level, Guinney et al. proposed four consensus molecular subtypes (CMSs) with different molecular and clinical features^4^. The TME of CRC consist of distinctive cell subpopulations, including tumor epithelial cells, cancer-associated fibroblasts (CAFs) and immune cells^5^. Stromal cell signatures detected from bulk transcriptomes were reported to be associated with risk of relapse and predict survival time of patients^6^. To explore the prognostic values of these multi-level biological features of CRC, Guinney et al. developed a multivariable Cox model for disease-free survival (DFS)^7^. They obtained the relative proportion of explained variation for clinicopathological features (tumor stage, age, sex and primary tumor location), CMS scores, genomics markers (MSI, BRAF and KRAS mutations), and the abundance of ten cell subtypes from bulk transcriptomes. The results showed that TME infiltration patterns could represent potent determinants of the recurrence risk for early-stage CRC. The constructed multivariable models revealed that the prognostic value of CMS and MSI could be explained by CytoLym and CAF infiltration patterns. The diversity of the TME also could be used for classification of cancers regarding prognosis^8^. Bruni. et al. summarized the prognostic significance of the major immune components in CRC ^8^. Galon et al. proposed an immunoscore system based on the density of CD3+, CD8+, or CD45RO+ lymphocytes obtained from immunohistochemistry (IHC) staining. They found that the immunoscore showed higher prognostic value than the TNM stage^9,10^. However, the c-indexes of this system for relapse-free survival (RFS) and overall survival (OS) were only 0.62 and 0.58 in their benchmarking cohorts^9^. There is still an urgent need for TME-related prognostic models with improved predictive accuracy.

Traditional transcriptomes have largely depended on analysis of bulk tissues, which obscures the signals of distinct cell subtypes. Single-cell RNA sequencing (scRNA-seq) provides unprecedentedly detailed characterization of the heterogeneity of cell transcriptomes, allowing the assessment of the complex tumor microenvironment^11,12^. scRNA-seq studies have been carried out in CRC^13–14^, and the main results of the applications of single-cell RNA technology in CRC have been summarized in **Table S1**. These studies provided valuable data source and could help in understanding the heterogeneity of TMEs of CRC. However, only a few cell types have been reported to be prognostic and few studies focused on developing TME-related prognostic models using scRNA-seq data. Application of scRNA-seq in translational research remains a challenge, partly due to the low library sizes, high noise and a large number of dropouts in scRNA-seq data, which might induce large experimental batch effects. Tan et al. developed a method to annotate cell types by comparing the expression of pairs of genes within each cell^17^. The relative expression orderings (REOs) of genes within a sample are robust against batch effects of experiments^18^ and could not be influenced by the normalization of datasets^19,20^, making them promising for developing prognostic models using bulk transcriptome data^21,22^. Wang et al. proposed single-cell pair-wise gene expression (scPAGE) to improve bulk RNA-seq data classification in acute myeloid leukemia^23^. The application of rank-based method in developing prognostic models from scRNA-seq data remains to be investigated. To the best of our knowledge, REO-based prognostic risk models from scRNA-seq in CRC have not yet been studied. In this study, we developed a data analysis framework to detect prognostic subpopulations and identify individualized recurrence risk signatures based on within-cell REOs of gene pairs to improve the risk stratification of patients with stage II–III CRC.

## Material and methods

### ScRNA-seq data preprocessing

We collected three scRNA-seq datasets from Gene Expression Omnibus (GEO) (**Table S2**). The Seurat package (https://satijalab.org/seurat/articles/get_started.html) was used to pre-process each scRNA expression profile based on three metrics including the expression frequency in cells per gene, number of measured genes and proportion of mitochondrial gene count per cell. To remove low-quality cells and doublets, we filtered out genes in less than three cells, removed cells with less than 200 or more than 6,000 genes, as well as cells with more than 20% mitochondrial gene expression in gene counts.

### Dimension reduction and annotation

The classic workflow in Seurat was used to perform dimension reduction and unsupervised clustering for each of the scRNA-seq datasets. Notably, the optimal value of parameter ‘resolution’ of *FindClusters* was determined by clustree^24^ package. To ensure that annotation results could be compared among different scRNA-seq datasets, we combined manual and supervised annotations (**Figure 1-Step1**). In the training dataset (GSE144735), we identified 7 major cell subpopulations according to the expression levels of classic markers. Then, we applied the same workflow for each major cell type to further reduce dimension and clustered cells into sub-populations based on several classic markers. As tumor-derived epithelial cells were heterogeneous in different patients, we used *classifyCMS* in CMSclassifier^25^ package to assign a CMS label to each tumor-derived epithelial cell. To apply the established landscapes of cell subtypes to two independent scRNA-seq datasets (GSE132465 and GSE132257), we used SciBet^26^, a supervised cell type annotation toolkit, to predict cell identities for cells from training set to query sets.

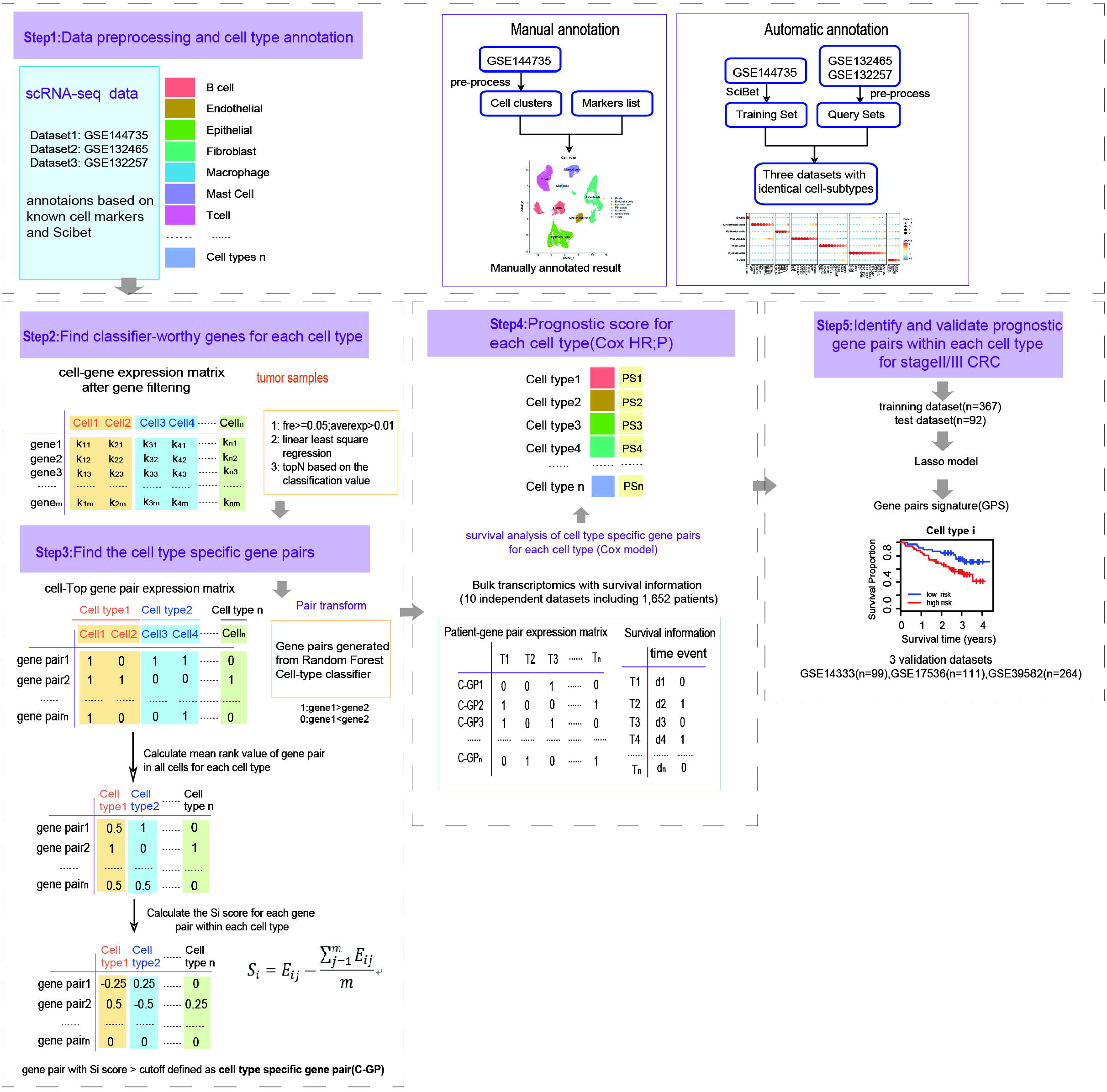

### Bulk transcriptomic data and preprocessing

We downloaded the CEL format files of ten bulk microarray datasets from GEO (**Table S2**). The expression values were normalized and log2 transformed. We further matched the probes with gene symbols in expression profiles according to the platform probes file, respectively. Probes matched to no or more than one gene were deleted. For a gene mapped to multiple probes, the arithmetic mean of the values of the multiple probes was calculated for its expression value. Finally, we downloaded clinical information files including TNM stages, gender, age, adjuvant chemotherapy and survival information (**Table S2**).

### Overview of scRank construction

#### i. Identification of cell type specific gene pairs with cell subpopulation-classifying value

Tumor-derived cells were used to identify cell subtype specific gene pairs. The *GetClassGenes* function from the singleCellNet^17^ package was used to select discriminative feature genes for each cell type according to the following criteria: detected in at least 5% of cells; the average expression value within cell type was larger than a pre-defined threshold (0.01 in this study); the classification value for cell type based on linear least square regression analysis ranked in top 100 genes (**Figure1A-Step2**). Then, we transformed the remaining data into a matrix from pairwise comparisons of the selected features for each cell. (**Figure1A-Step3**). The linear least square regression analysis was performed to select gene pairs with top 1000 ranked by the classification value for cell types. Based on this filtered gene pair expression matrix, we used function *scn_train* to train a random forest (RF) classifier for each cell subtype. The AUPR (Area Under the Precision-Recall curve) was used to assess the performance of the classifiers. For gene pairs in RF classifier of cell subtypes, we calculated the specific-score *S^i^* for each gene pair in all cell subtypes as follows.

Firstly, we calculated the average REO for each gene pair in cell subtype j:

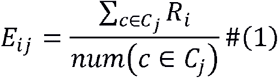

where *C_j_, i, c* and *R_i_* represent the cell subtype j, gene pair *i*, cell *c* and the REO of *i* in *c*, respectively. Additionally, if gene pair *i* was not cell subtype specific, the value of *E_ij_* in all cell subtypes should be similar. In other words, the *E_ij_* in one cell subtype approximated to the mean of *E_ij_* in all cell subtypes:

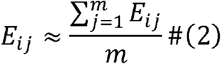

The m represented the number of cell subtypes.

If *E_ij_* was higher in a cell subtype compared with other cell subtypes and the difference value was greater than a cutoff, we considered it was a specific gene pair. To better understand each gene pair’s specificity, we calculated specific-score *S_i_* for each gene pair per cell subtype:

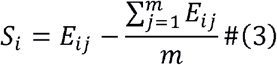

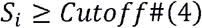

When *S_i_* was greater than Cutoff (0.6 in this study), we regarded pair *i* as the cell subtype specific gene pair (C-GP) for cell subtype j.

#### ii. Evaluation of prognostic value for each cell type

Let Ga and Gb represented the expression values of gene a and gene b, respectively. We applied the univariate Cox proportional-hazards regression model to evaluate the correlation of the REO pattern (Ga >Gb or Ga < Gb) of each C-GP with the recurrence survival time of CRC patients. P value was adjusted by the Benjamini and Hochberg (BH) method. Furthermore, C-GPs with adjusted P value less than 0.2 were defined as recurrence-related. In addition, we applied the C-GPs of T cells, Epithelial cells, Endothelial cells and Fibroblasts on a bulk RNA-seq with fluorescence-activated cell sorting (GSE39396)^27^ to evaluate their cross-platform stability.

#### iii Development of individualized recurrence risk signatures

To identify individualized recurrence risk signatures for each cell subtype, we aggregated C-GPs from three scRNA-seq datasets:

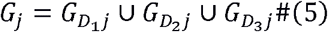

*D_x_, j* represented the scRNA-seq dataset x and cell subtype j, respectively. *G_D_x_j_* represented all the C-GPs of cell subtype j in scRNA-seq dataset x. In addition, we collected seven bulk RNA-seq datasets (n=459) and 80% of which were randomly sampled as the training dataset while the rest samples were used as the test dataset. The other three independent bulk RNA-seq datasets GSE14333 (n=99), GSE17536 (n=111) and GSE39582 (n=264) were used as the validation datasets (**Table S2**). We applied the lasso-cox model ^28^ to train a signature which was then used to score the recurrence-risk for each patient. We chose ‘lambda. 1se’ as the final penalty for the model and *surv_cutpoint* was used to work out the optimal cutoff of the risk value. We calculated the risk-score based on the REOs of C-GPs and corresponding coefficients for each patient. A sample was determined to be high risk if the corresponding risk score was higher than an optimal cutoff, otherwise, this sample was labeled as the low-risk. After that, we performed Kaplan-Meier analysis using the *survdiff* function to test the difference of recurrence survival time between two risk groups. Finally, the multivariable Cox proportional-hazards regression model was used to evaluate whether signature performed as an independent prognostic factor after adjusting for other clinical factors including tumor stage, gender, age, primary tumor location, and gene mutations (BRAF, KRAS and TP53) and mismatch repair status.

### Determining cell type abundance from bulk transcriptome

Based on the HVGs-cell expression matrix for each scRNA-seq dataset, we inferred the abundance of each cell subtype in bulk transcriptome profile from the signature matrix established by CIBERSORTx^29^. Similarly, we applied cox proportional hazard model to evaluate the contribution of the infiltration of each cell subtype to the recurrence survival time.

### Prognostic signature enrichment analysis

We used *coxph* to generate the risk-associated (HR>1) and protected-associated (HR<1) gene sets in each bulk transcriptome dataset (FDR < 0.2). Then, we averaged the genes’ expression of all cells in each cell subtype in the scRNA-seq dataset. Single sample gene set enrichment analysis^30^ (ssGSEA) was performed to calculate enrichment scores with prognostic-associated gene sets.

### Identifying prognostic-associated subpopulations using Scissor

Scissor^31^, a module to distinguish clinically phenotype relevant cells in scRNA-seq dataset, was used to calculate the pertinence relation between single cell in scRNA-seq profile and single sample in bulk RNA-seq profile. We used default parameters cox mode of Scissor to detect prognostic-associated cells in each scRNA-seq profile.

### Evaluation of performance for methods used for prioritizing prognostic-associated subpopulations

Besides the methods mentioned above, we used *FindMarkers* function from Seurat package to detect the genes differentially expressed among cell subtypes. We named this method as Uni-Markers. Similarly, we applied the *coxph* function to calculate the HR of each differentially expressed gene in cell subtypes for each scRNA-seq dataset. To compare the performance of different methods (CIBERSORTx, GSEA, Scissor, Uni-Markers and scRank), we performed a literature review on the prognostic value of several cell subtypes (**Table S3**). We used ‘R’ (risk) and ‘P’ (protected) to label the prognosis of each cell subtype. Under this circumstance, if the current result was consistent with the previous study, we announced it positive (‘p’ as a proxy) for the prognostic contribution. If not, it was labeled negative (or ‘N’). Besides, ‘A’ (ambiguous) referred the prognostic value had conflict results in different scRNA-seq datasets or even cannot obtain the prognostic classification. ‘U’ (uncertain) represented the prognostic value of one cell subtype was unclear in prior research. Further, we calculated three quantitative metrics to assess performance of these methods:

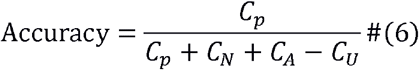

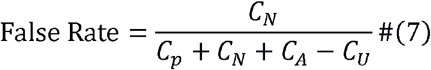

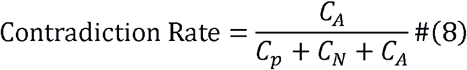

*C_p_* represented the number of cell-subtypes whose prognostic contribution was in line with previous study and different scRNA-seq datasets, while *C_N_* represented the number of cell subtypes with similar prognostic-predict among different scRNA-seq datasets but were inconsistent with previous study. *C_A_* and *C_U_* represented the number of cell subtypes of which prognostic contribution were evaluated as ‘A’ and ‘U’, respectively.

## Results

### Cell identity annotation across independent datasets

Firstly, we manually annotated the major cell types of a scRNA-seq dataset (GSE144735) (**Figure 2**). Based on the expression of classic markers (**Figure 2B, Figure S1B-C**), 4 immune cell types (T cells, B cells, Myeloid cells and Mast cells) and 3 non-immune cell subtypes (Endothelial cells, Fibroblasts and Epithelial cells) were distinguished. Then, we used SciBet^26^ to find the discriminative feature genes (**Figure S1A**) in the training dataset via E-test. After that, we annotated two independent scRNA-seq datasets (GSE132465 and GSE132257) separately (**Figure S1A-B**). Seven major cell types similar to the training dataset were obtained and UMAP of classic markers’ expression level for each cell type were also shown (**Figure 2B, S1A-B**). Next, we reapplied dimension reduction and clustered each major cell types into minor subtypes. For T cells, we identified 10 subpopulations: Naïve T cells, CD8+ T cells, CD4+ T cells and others. Furthermore, CD8+ T cells were separated into Tex (Exhausted T cells) and CTLs (Cytotoxic T cells), which were named as GZMK+ CTL and KLRD1+ CTL. CD4+ T cells were separated into four subtypes, including CD4+ Tfhs (Follicular T-helper cells) and Treg (regulatory T cells). Other T cells were separated to γδ T cells (Gamma-delta T cells) and NK cells (Natural killing cells) (**Figure S2A-B**). We also identified seven sub-populations of B cells (**Figure S3A**). In particular, there are two distinct IgA+ plasma B cells clusters (**Figure S3B**). One cluster highly expressed IGLC2 and the other highly expressed IGLL5. Note that the IGLC+ IgA+ plasma B cells cluster was also identified in CRC by Wang et al.^32^. For Myeloid cells, we identified Dendritic cells (DCs) and Monocyte-Macrophages.

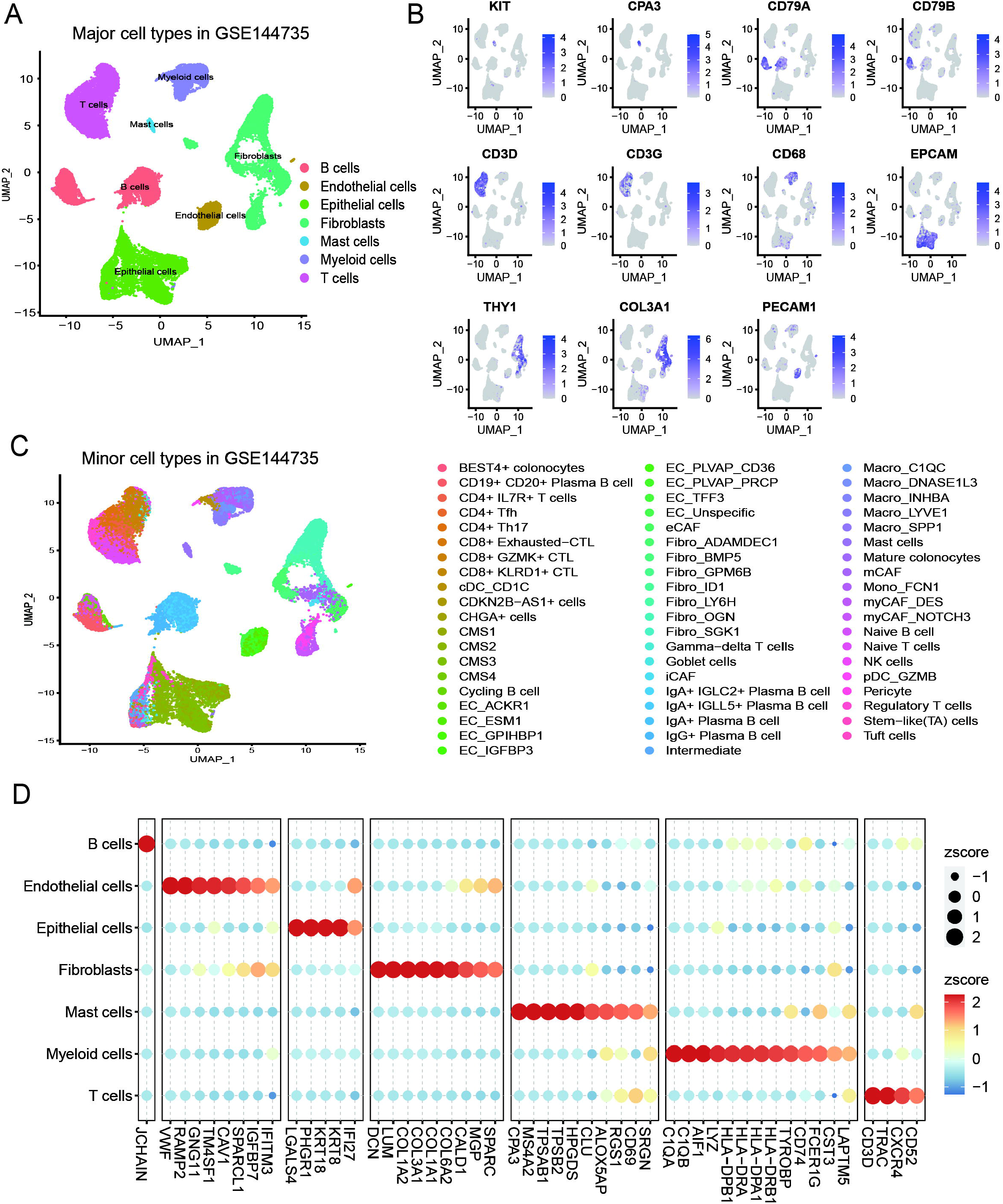

For non-immune cell types, we identified CAFs, eCAF (Extracellular CAF) and myCAF (myofibroblast), which were named based on differential expressed makers^14,33,34^ (**Figure S5**). Using the markers from A. Sharma^35^, we identified 8 sub-populations of endothelial cells (**Figure S6**). We found that the distribution of epithelial cells was heterogeneous among different patients (**Figure S7C**), therefore we separated epithelial cells from tissues. For epithelial cells from normal tissues, we applied classic markers^14^ to identify subpopulations (**Figure S7A**). We assigned CMS labels to epithelial cells from tumors and nearby tissues by calculating the enrichment score of the CMS-related signaling pathway gene sets. It turns out that CMS4-like cells enriched genes related to EMT and TGFβ signaling pathways. CMS2-like cells mainly enriched genes related to WNT and MYC signaling pathways. CMS3-like and CMS1-like cells enriched genes related to differentiation and MSI, respectively (**Figure S7B**). These results were consistent with Lee et. al^14^ and in accordance with the conventional understanding of CMS molecular subtypes^4,36,37^.

Cell sub-populations were also identified in GSE132465 and GSE132257 datasets using the same workflow (**Figure S8)**. The proportions of cell sub-populations were shown in **Figure S9**. As for Mast cells with low cell counts, no further subtypes were annotated.

### Prioritizing prognostic-associated subpopulations based on cell type specific gene pairs

Firstly, we identified cell type specific gene pairs (C-GPs)(**Figure 1 Step 2-3**). SingleCellNet^17^ was used to train RF classifiers for each cell subtype based on gene pairs in the three independent scRNA-seq datasets, respectively. Then, we plotted the Precision-Recall (PR) curve and calculated AUPR to evaluate the performance of the classifiers. In general, the classifiers performed well for most cell subtypes (**Figure S10**), suggesting that cell subtypes could be annotated accurately based on REOs. We selected C-GPs for each cell type and observed that cell sub-populations derived from the same major cell type shared some specific gene pairs as expected (**Figure 3A, S11A**). In addition, these cell sub-populations still had their own specific gene pairs (**Figure 3A, S11A**). For example, we randomly selected 20 IgA+ Plasma B cells in all scRNA-seq datasets and plotted the REOs of B2M-IGKC, respectively. It was found that this gene pair maintained cross-dataset stability (**Figure 3B**). Another example was the specific gene pair CCL5-GADD45B of CD8+ KLRD1+ T cells. Notably, we observed that the REOs of this gene pair were consistent (**Figure 3B**).

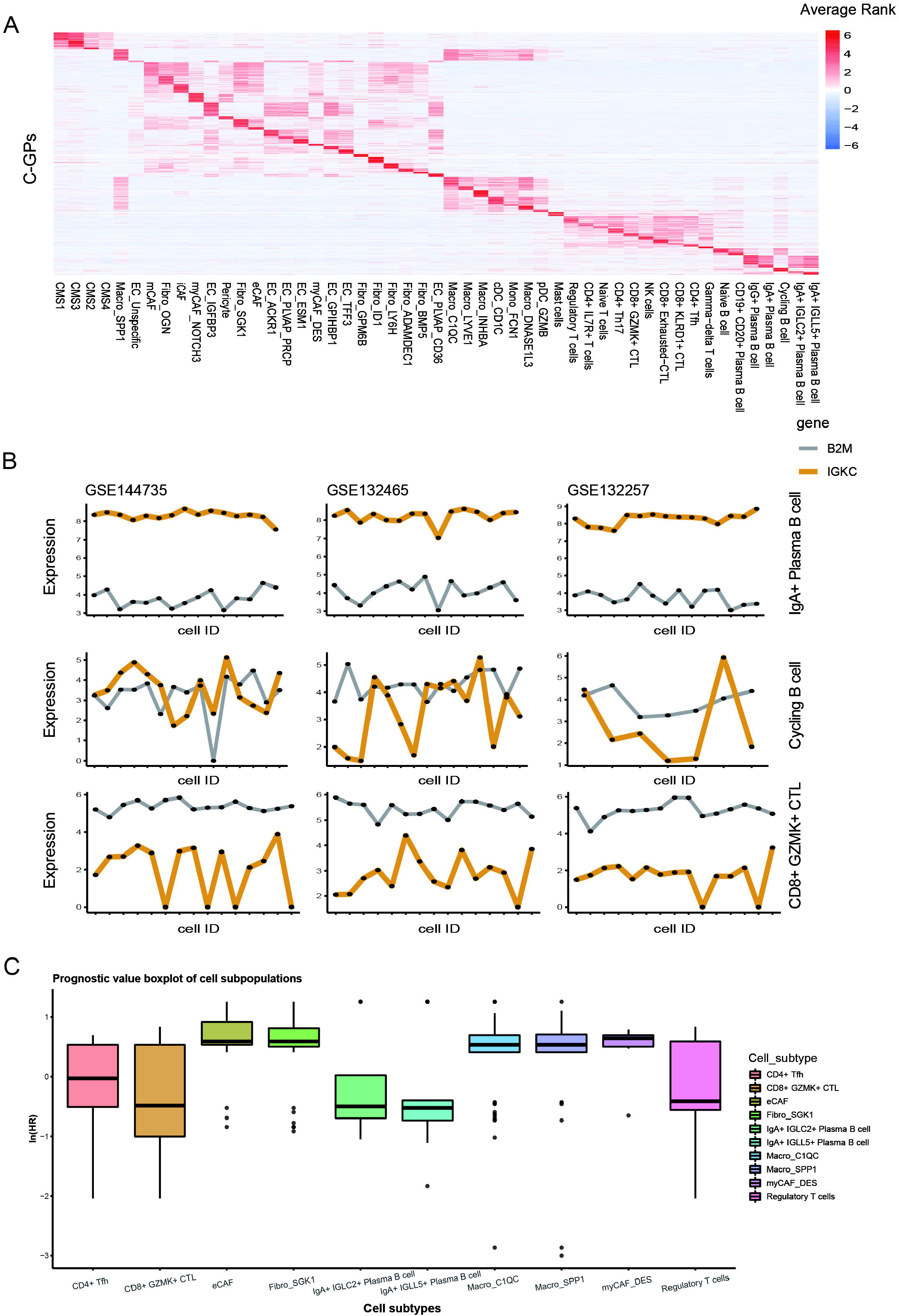

Next, for each scRNA-seq dataset, we applied univariable cox model to correlate the C-GPs and DFS of patients. The prognostic values of several cell types evaluated here were in line with previous studies (**Table S3**). T cells’ sub-populations, such as CD4+ Tfh and CD8+ GZMK+ CTL correlated to good prognosis (**Figure 3C**). CD4+ Tfh was shown to be an independent prognostic predictor in breast cancer and correlated to improved prognosis^38^. CD8+ GZMK+ CTL, a significant component of cell-mediated immunity, plays a central role in tumor cytotoxicity^39^. Besides, the results of other cell subpopulations were consistent with previous studies (**Figure 3C**). For example, in B cells’ sub-populations, low-expressed IGLC2 was considered as a factor correlated to worse prognosis in Triple Negative Breast Cancer patients^40^ and the expression level of IGLL5 was positively correlated to tumor size in clear cell renal cell carcinoma^41^. When it comes to the sub-populations of Myeloid cells, it has been reported previously that Macro_SPP1 and Macro_C1QC correlated to worse prognosis for CRC patients^15^. As for sub-populations of Fibroblasts, eCAF that highly expressed CST1, was reported to correlate to worse-prognosis and tumor generating in CRC^42^. myCAF was found to promote cancer development and progression^43^.

### C-GPs achieves higher precision compared with current methods

The performance of C-GPs was compared with four methods in common practice. Firstly, we used CIBERSORTx to evaluate the relevance between cell subtype infiltration and prognosis of CRC patients from bulk transcriptome, which was performed in most existing studies^44–46^. We found that the infiltration of a few cell subtypes, such as eCAF and EC_GPIHBP1, correlated with worse prognosis in CRC patients (**Figure S12**). Fibro_ADAMDEC1 and EC_IGFBP3 were related to good prognosis (**Figure S12**). Secondly, the results of ssGSEA revealed that prognosis-protected genes mainly enriched on the sub-populations of B cells and T cells (**Figure S13**). While the recurrence-related genes mainly enriched on fibroblasts, myeloid cells and endothelial cells, especially SPP1+ and C1QC+ macrophages (**Figure S14**). Thirdly, Scissor was used to integrate the phenotype of patients in bulk transcriptome data and cells in scRNA-seq data. With Scissor, we observed that Macro_SPP1, Macro_INHBA, Fibro_OGN, Fibro_SGK1, CAFs, CMS1-like cells and CMS3-like epithelial cells were related to CRC recurrence. CMS2-like epithelial cells were related to good prognosis. Interestingly, results of subtypes of B cells and T cells from Scissor were not associated to clinical outcome, which may indicate potential limitations of this method (**Figure S15**). Finally, applying Uni-Markers, many cell subtypes including CD8+ GZMK+ CTL and cDC_CD1C were recognized as related to prolong patient’s DFS time. However, it could not evaluate exactly for CAFs and some macrophages, which were considered as risk-factors in CRC ^6,47–50^.

To have a more systematic evaluation on these methods, we used three metrics (Accuracy, False Rate and Contradictory ratio, see Method) to compare the prognostic value of these cell subtypes with consensus results from existing literature (**Table S3**). C-GPs showed high accuracy (Accuracy rank: C-GPS > GSEA > CIBERSORTx > Uni-Marker > Scissor), low false rate (False rate rank: C-GPS < GSEA ≈ CIBERSORTx ≈ Scissor < Uni-Marker) and low contradictory ratio (GSEA < C-GPS < CIBERSORTx < Uni-Marker ≈ Scissor) (**Figure 4**). Moreover, C-GPs ensure the stability cross different datasets and REO is a reliable signature for specific cell phenotypes in bulk RNA (**Figure S16**).

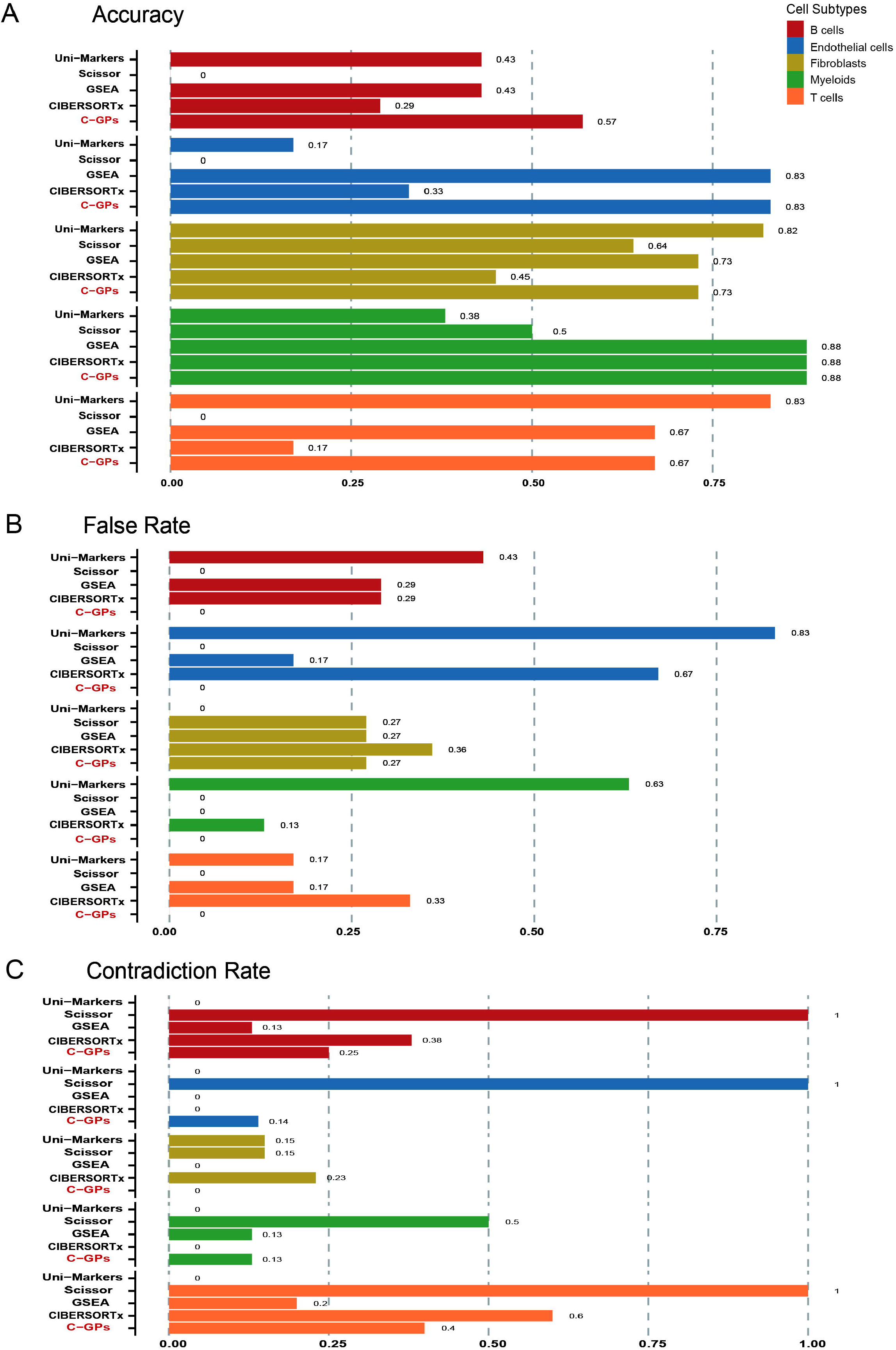

### Within-cell REOs of gene pairs predict recurrence risk in CRC

We used the C-GPs as features of each cell sub-population and used lasso-cox model to train a signature to predict the recurrence risk (**Figure 1-Step5**). Results show that the trained signature for 28 cell subtypes, including CD4+ Tfh and CD4+ IL7R+ T cell, were able to predict the recurrence risk for early stage(II/III) of CRC patients (**Figure 5**). The infiltration of CD4+ Tfhs was considered as a potent prognosis-predictor in breast cancer^38^. The expression level of IL7R was relevant to prolonged DFS and OS in Lung adenocarcinoma (LUAD)^51^. Besides, for non-immune cell sub-populations, the low-expression level of BPM5 was considered as a predictor to worse prognosis in CRC^52^. To further validate the signature derived by C-GPs, we selected Fibro_SGK1 as an example. The signature of Fibro_SGK1 consisted of 5 gene pairs (**Table S4**) and it performed well in training set (n = 367, P value < 0.001, log rank test; C-index = 0.68) and inner test set (n = 92, P value = 0.011, log rank test). Moreover, it also could separate patients into high-risk and low-risk groups exactly in three independent validation sets GSE14333 (n = 99, P value = 0.0032, log rank test), GSE17536 (n = 111, P value = 0.014, log rank test) and GSE39582 (n = 264, P value = 0.0053, log rank test) (**Figure 6A-B**). The key parameters of the lasso-cox model were shown (**Figure S17**). Furthermore, multivariable cox analysis on our signature and other clinical factors including age, gender, clinical stage and genomic biomarkers showed that our signature was still an independent recurrence-risk predictor (**Figure 6C**). Features and their weights were shown (**Table S4**). Similar results were observed in CD8+ GZMK+ CTLs and IgA+ IGLC2+ B cells (**Figure S18, S19**). Patients with high level SGK1 had shorter DFS and OS than those with low expression level SGK1 in esophageal squamous cell carcinoma^53^. CD8+ GZMK+ CTLs were related to the prolonged DFS^54^. IgA+ IGLC2+ plasma B cells were related to worse prognosis in CRC^32^. In general, CD8 CTLs have been extensively studied in the literature, while IgA+ IGLC2+ B cell and Fibro_SGK1 are novel cell subtypes which could potently predict the recurrence-risk in CRC.

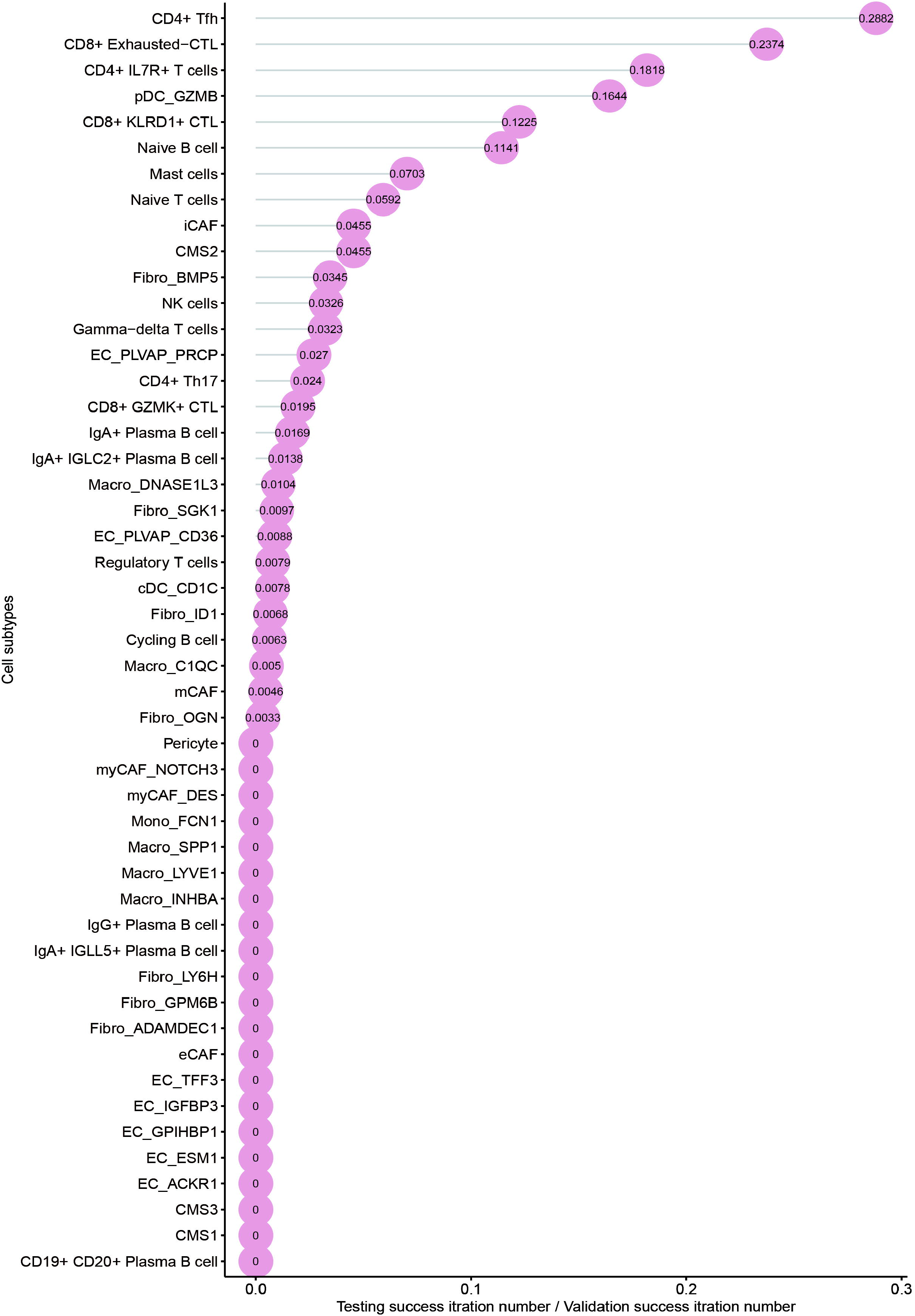

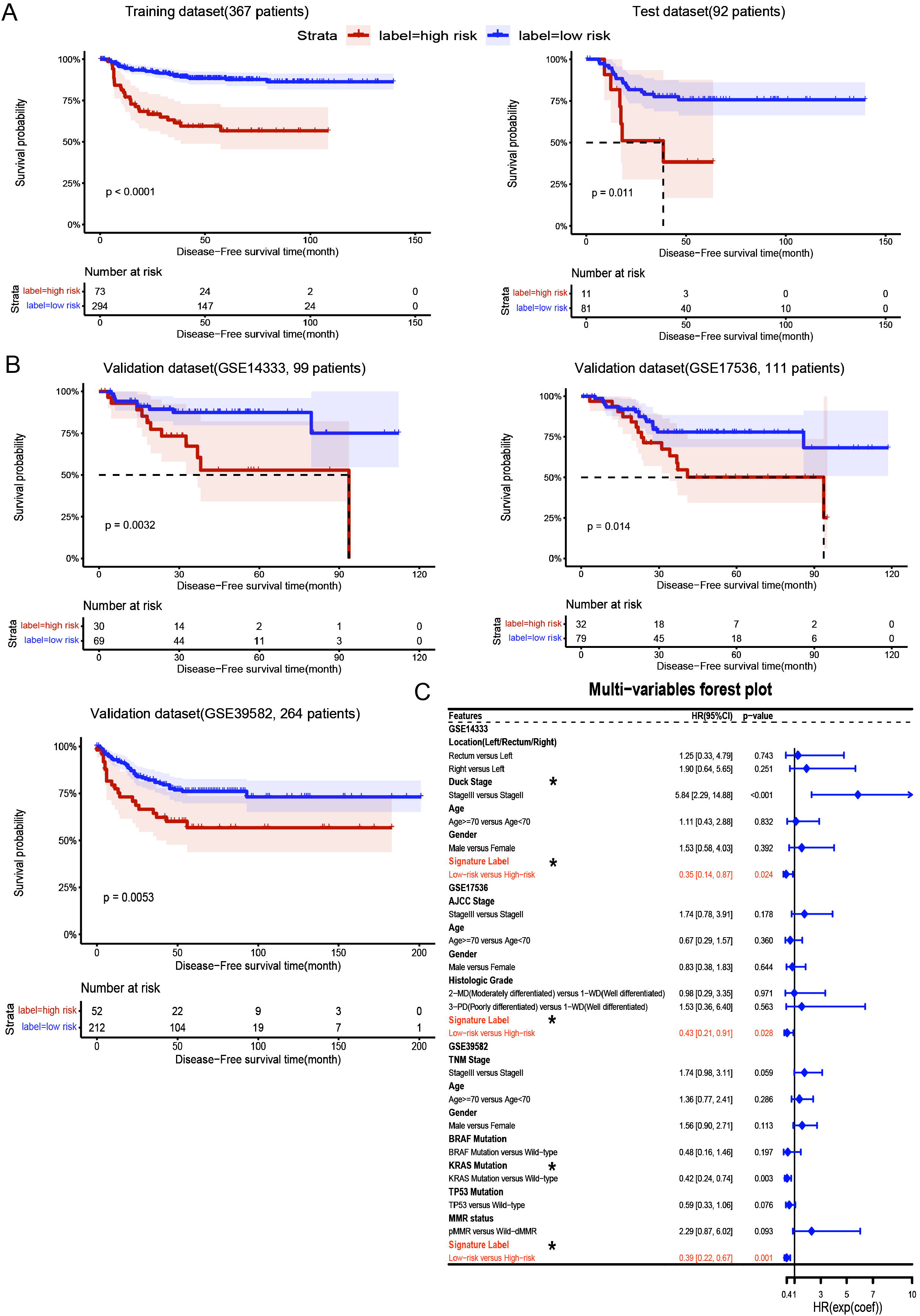

## Discussion

Recent advances in high-throughput technologies have facilitated the application of molecular biomarkers for prognosis prediction of CRC. However, most of the reported bulk transcriptional biomarkers were based on expression levels of the signature genes^55–57^. Due to experimental batch effects ^58^, the risk classification depends on the data normalization, which could not be diagnosed at the individualized level^21^. In contrast, the REOs of genes within a sample are robust against experimental batch effects and normalization methods^18,19^. It is necessary to screen REO-based signatures as prognostic biomarkers.

While TME has been reported to be important for tumor treatment response and survival of patients^8,59^, bulk tumor-based transcriptome only provides averaged data and could not accurately characterize the gene expression of subpopulations of the TME. The use of scRNA-seq in cancer research has improved our understanding of TME^60,61^. Identifying cell subpopulations that associated with clinical outcome could facilitate discovery of cell type targeted therapies as well as prognostic biomarkers. Most scRNA-seq datasets include fewer than 20 samples, which could not be used to identify the cell subpopulations associated with survival time for the lack of statistical power. Therefore, it is necessary to take full use of valuable clinical information to prioritize cell subpopulations from single-cell transcriptomes. In our study, we developed a workflow to identify cell subpopulations with specific REOs across datasets. Multiple variable COX model and Kaplan-Meier curve from our results confirmed the established role of cytotoxic T cells, which was related to prolonged disease-free survival time of CRC patients. In addition, we further identified two new cell subpopulations: IgA+ IGLC2+ B cells and Fibro_SGK1, which had strong relation to CRC recurrence. It should be noted that the REOs relied on accurate annotations of cell types. In addition, REOs could be affected by the dropout events in scRNA sequencing.

In conclusion, scRank^XMBD^ is a user-friendly toolkit to rapidly distinguish a set of gene pairs which could represent the property of cell subpopulation well. Furthermore, it could help to evaluate whether a novel cell subtype associates with survival when bulk RNA-seq data was available.

## Data availability

All scRNA-seq and bulk transcriptomes analyzed in this work were available in GEO database and summarized in **Table S2**.

## Author contributions

R.Y, J.H. and M.T. supervised the study. M.T., Y.L. and J.S. developed the framework and performed the analysis. M.T., YL., W.Y, J.S., Z.Z, J.X, S.L and C.L wrote and revised the manuscript.

## Key Points

- REOs-based signatures could classify most cell subtypes accurately in scRNA-seq data.
- Cell type specific gene pairs (C-GPs) were identified to evaluate prognostic value for each cell subpopulation. C-GPs achieves higher precision compared with four current methods.
- We developed single-cell gene pair signatures to predict recurrence risk for patients individually.
- A user-friendly toolkit, scRank^XMBD^ (https://github.com/xmuyulab/scRank-XMBD), was developed to facilitate the application of the rank-based method in scRNA-seq data for prognostic biomarker discovery and precision oncology.

