## Supplementary figures and images for "Prioritizing prognostic-associated subpopulations and individualized recurrence risk signatures from single-cell transcriptomes of colorectal cancer"

### Supplemental Figure1

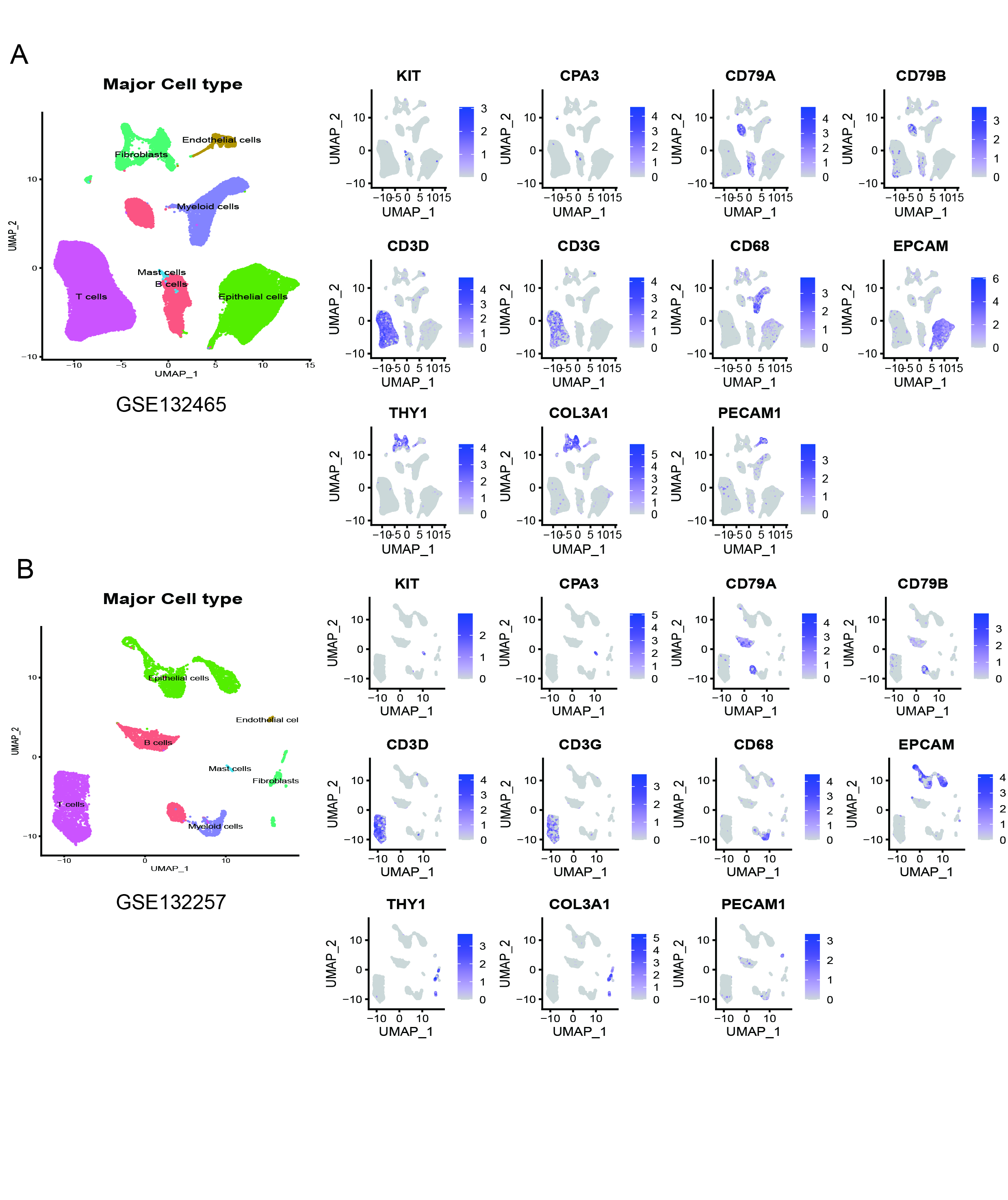

### Supplemental Figure2

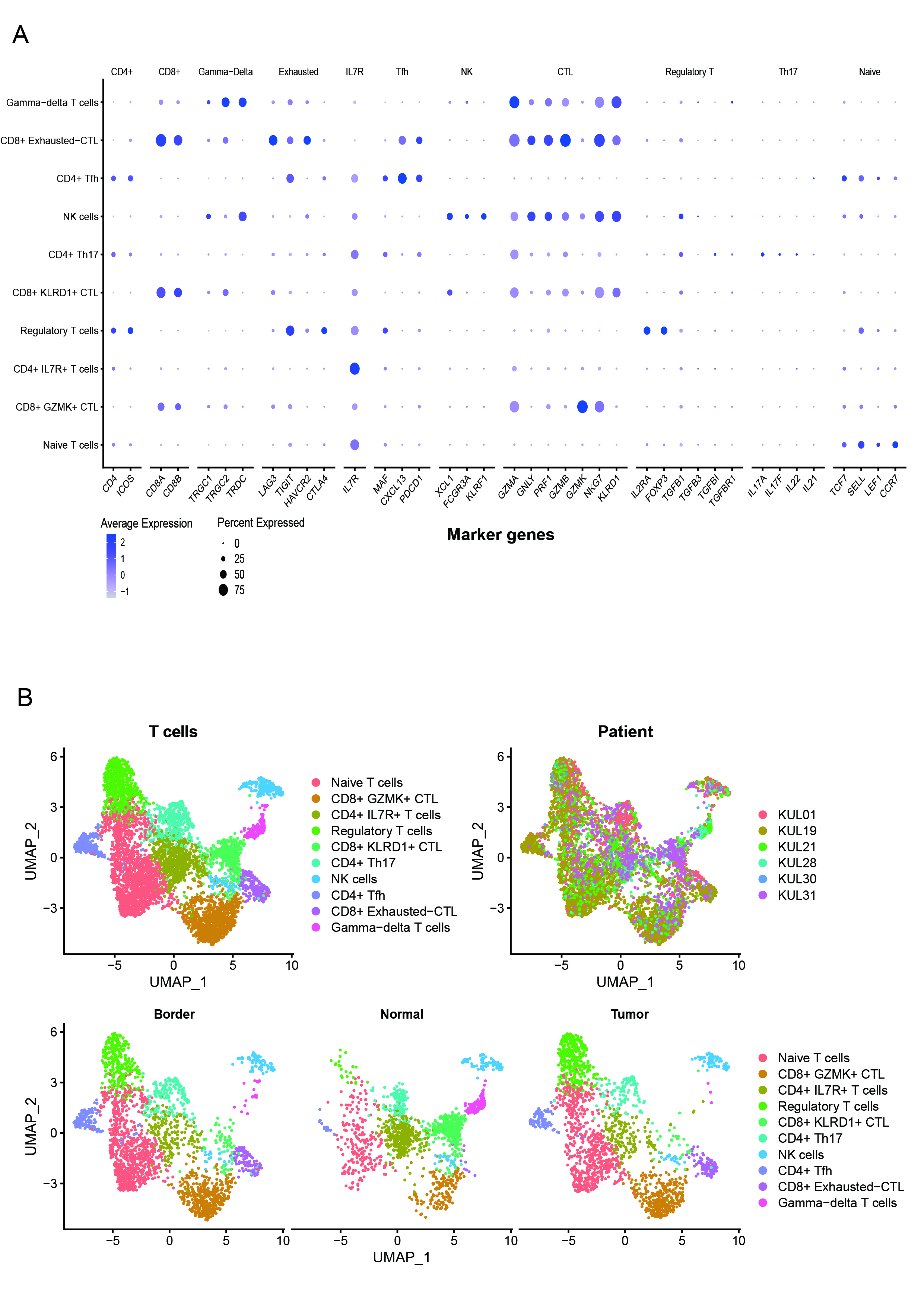

### Supplemental Figure3

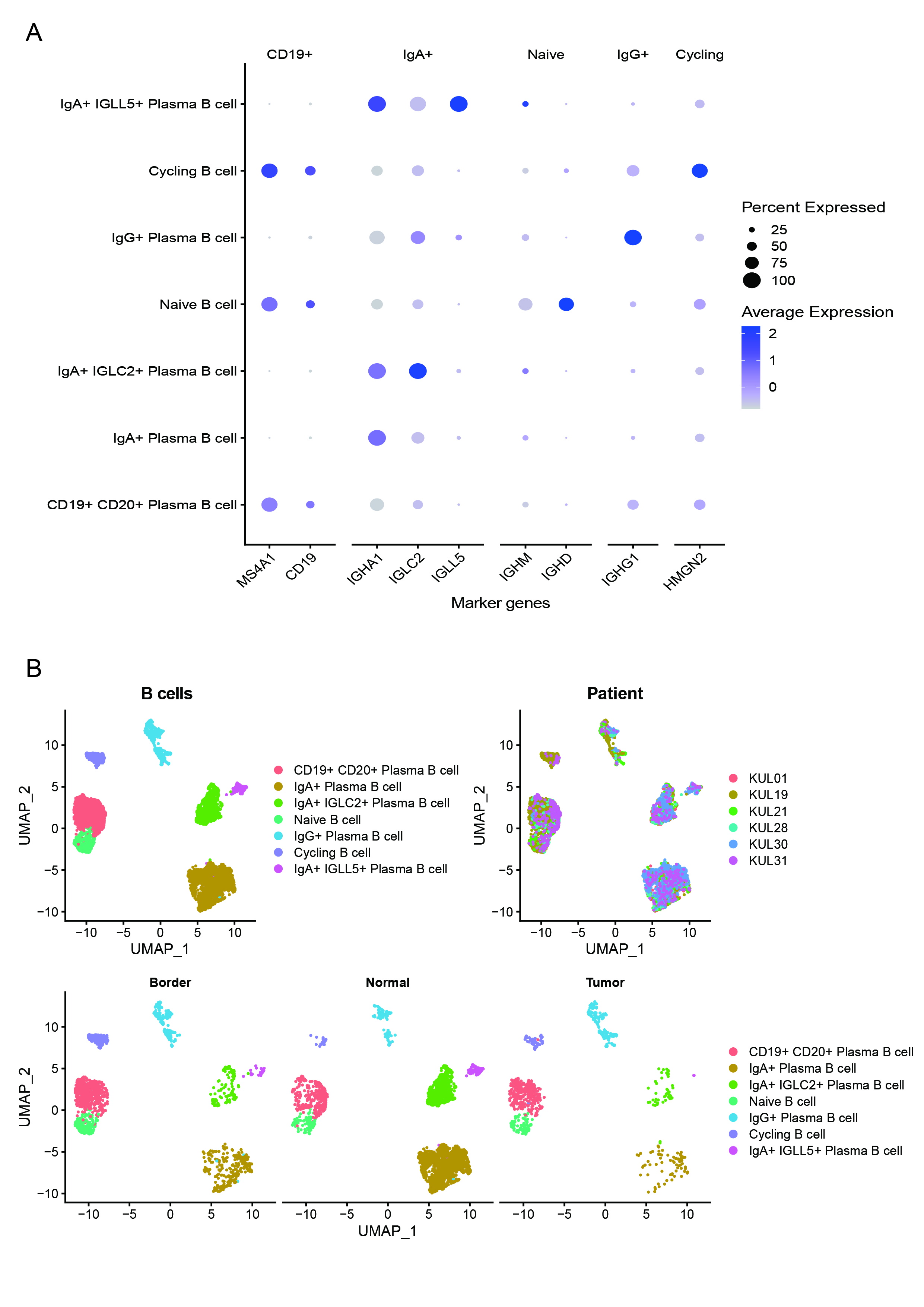

### Supplemental Figure4

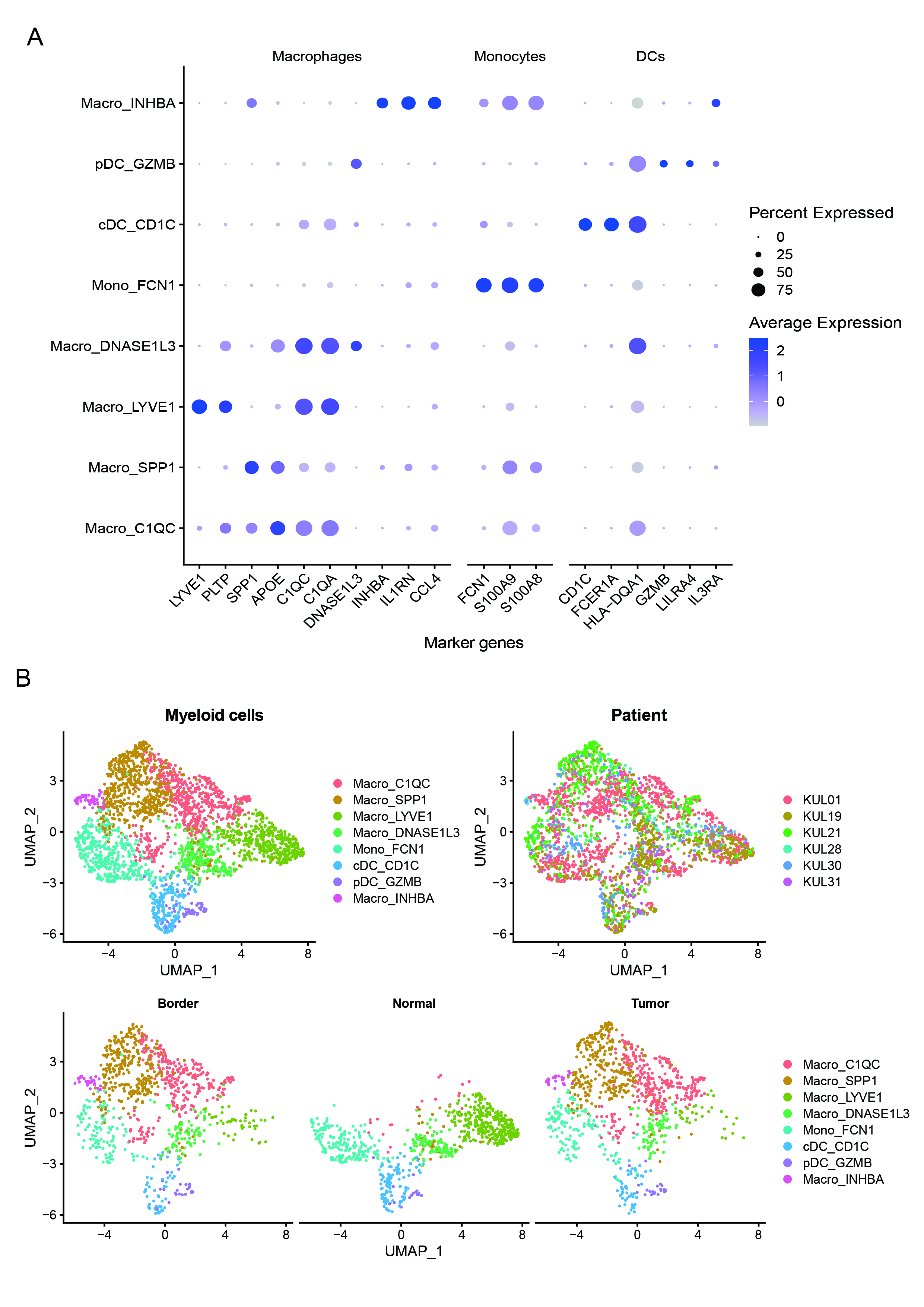

### Supplemental Figure5

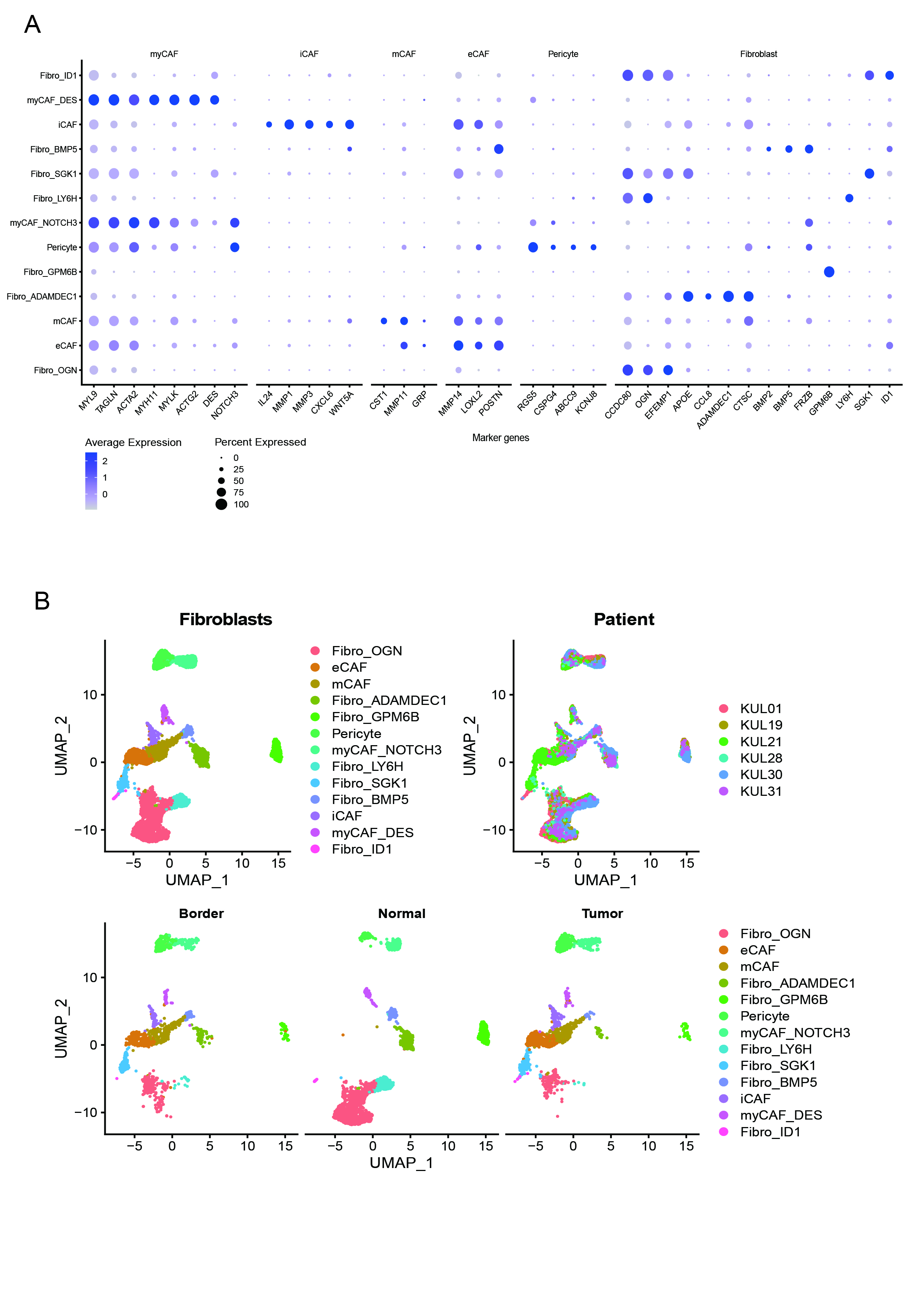

### Supplemental Figure6

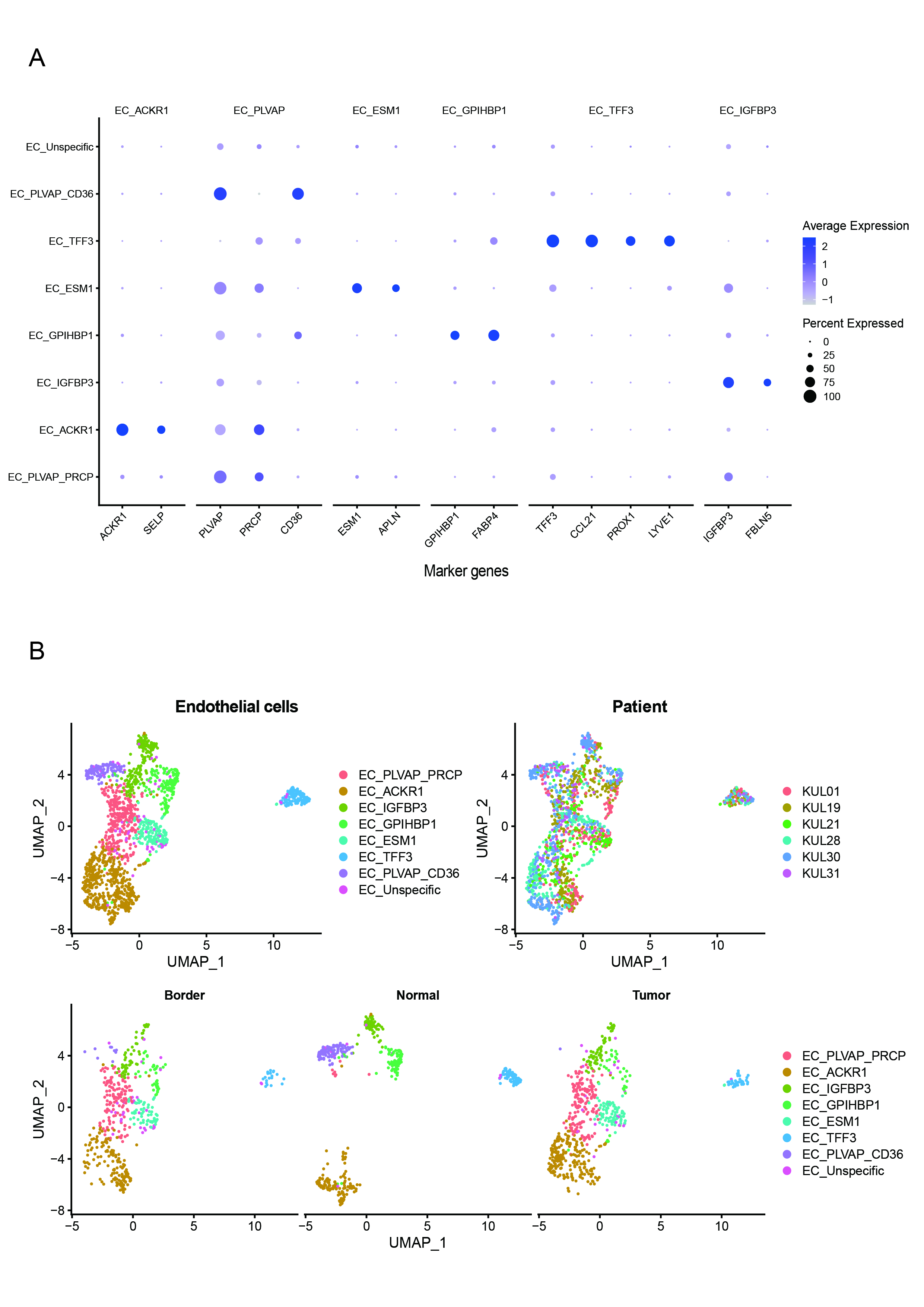

### Supplemental Figure7

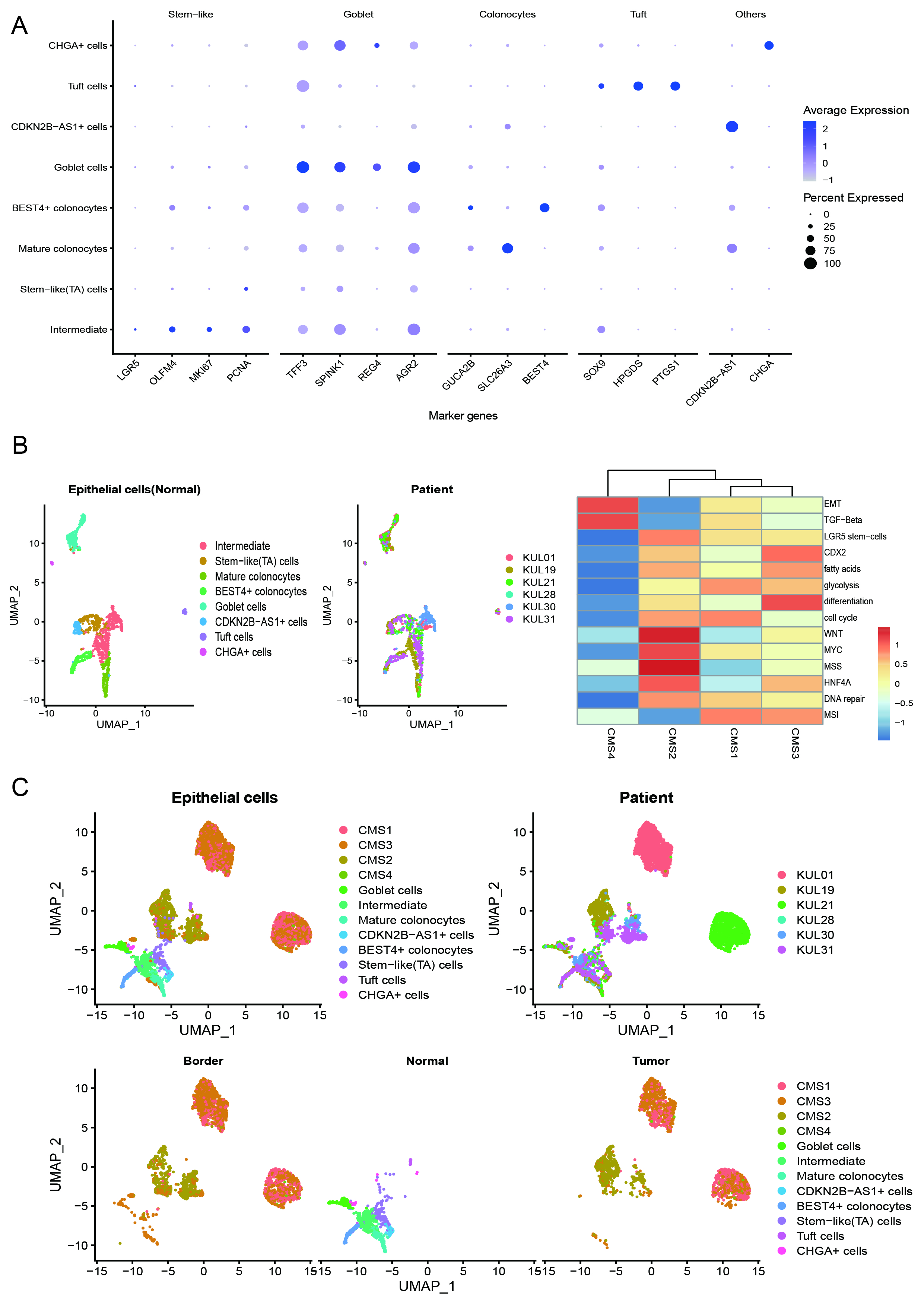

### Supplemental Figure8

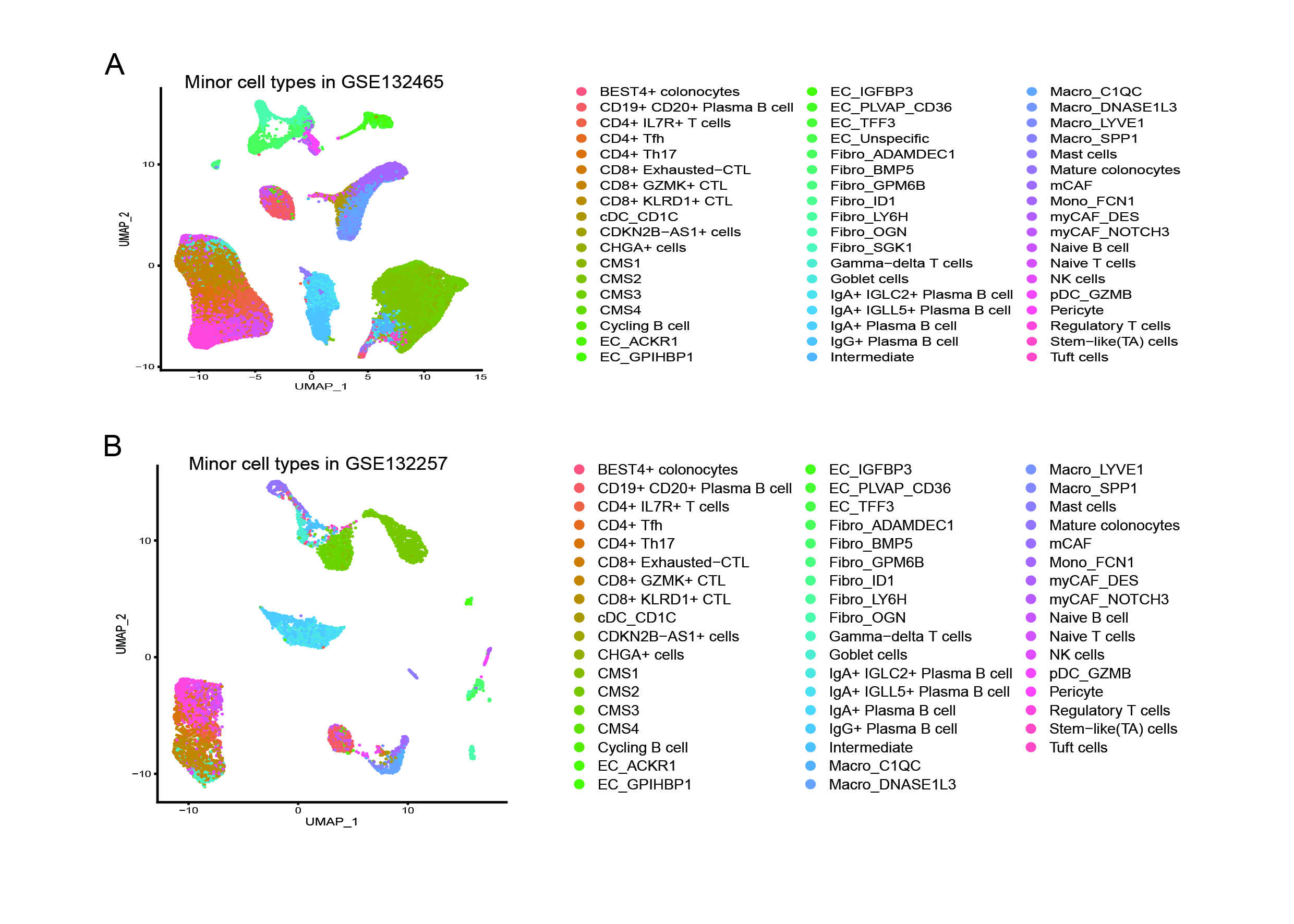

### Supplemental Figure9

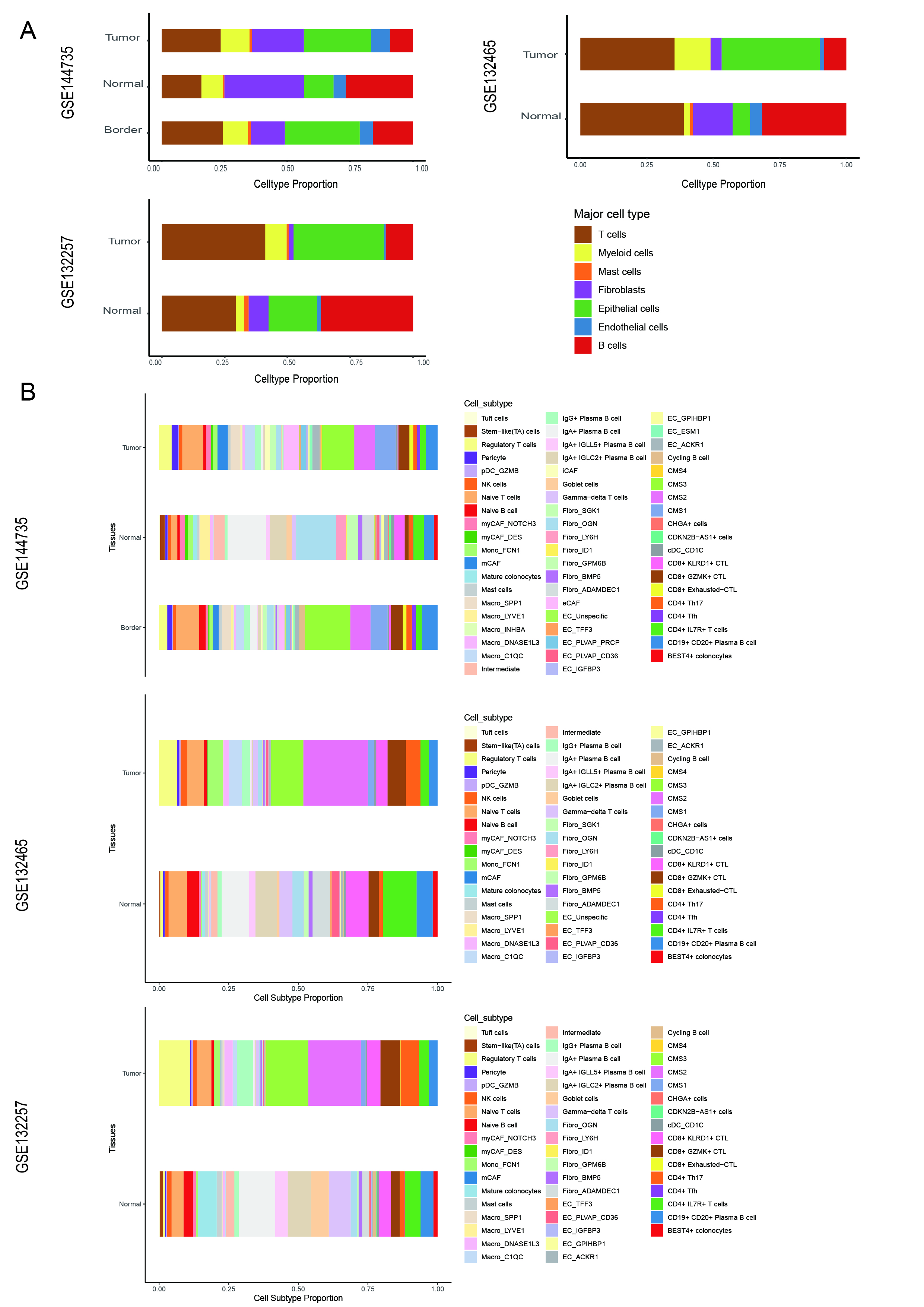

### Supplemental Figure10

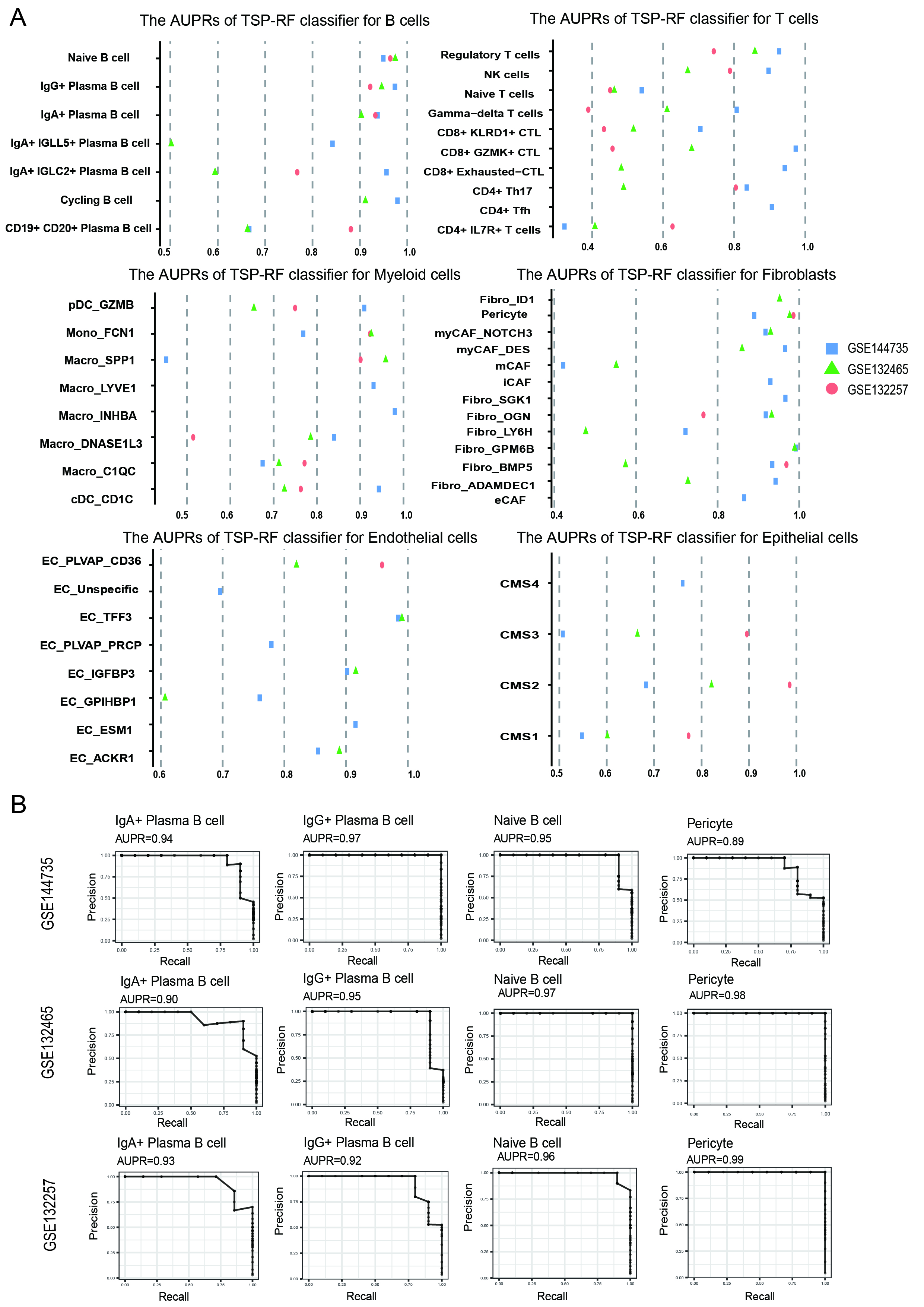

### Supplemental Figure11

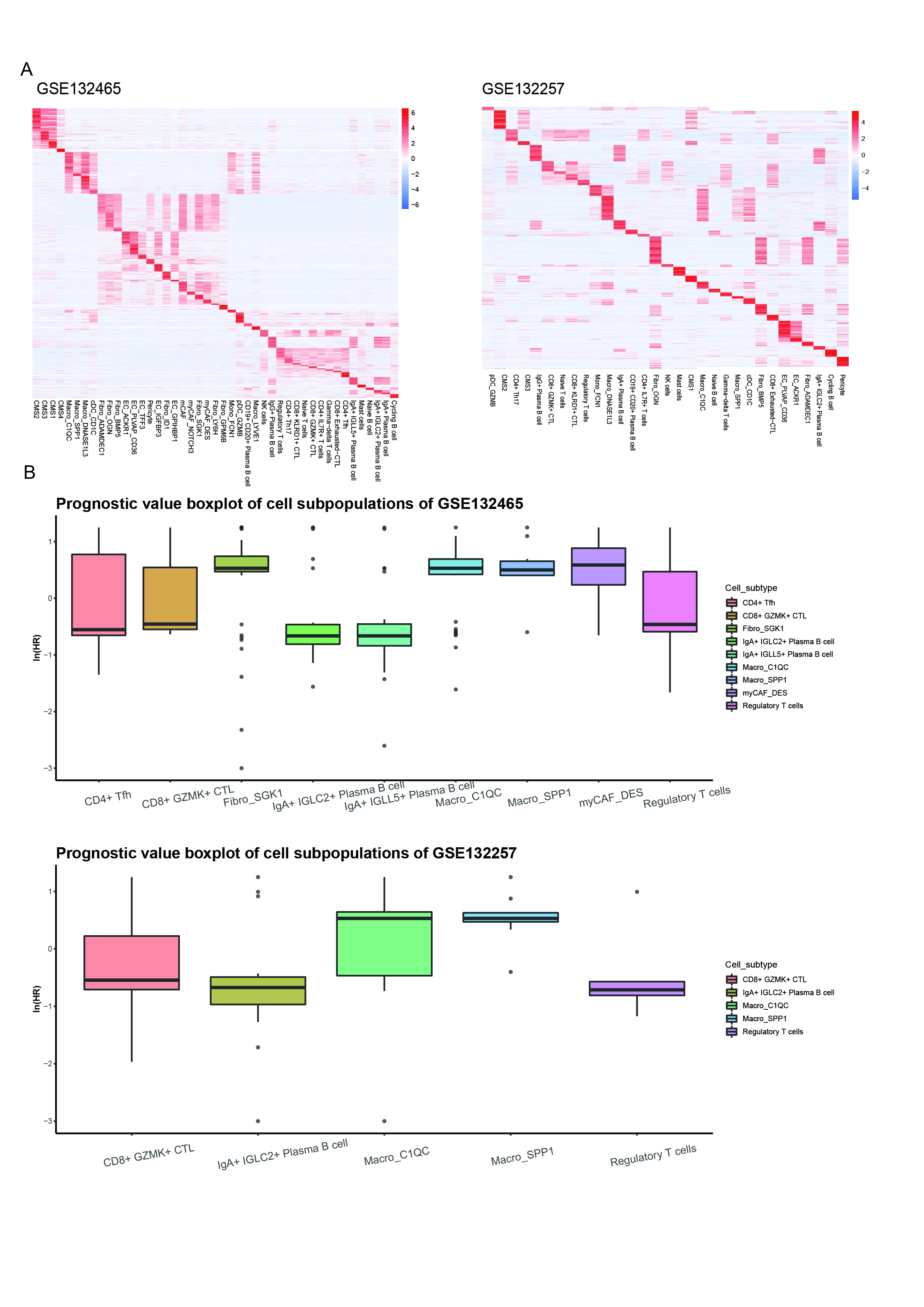

### Supplemental Figure12

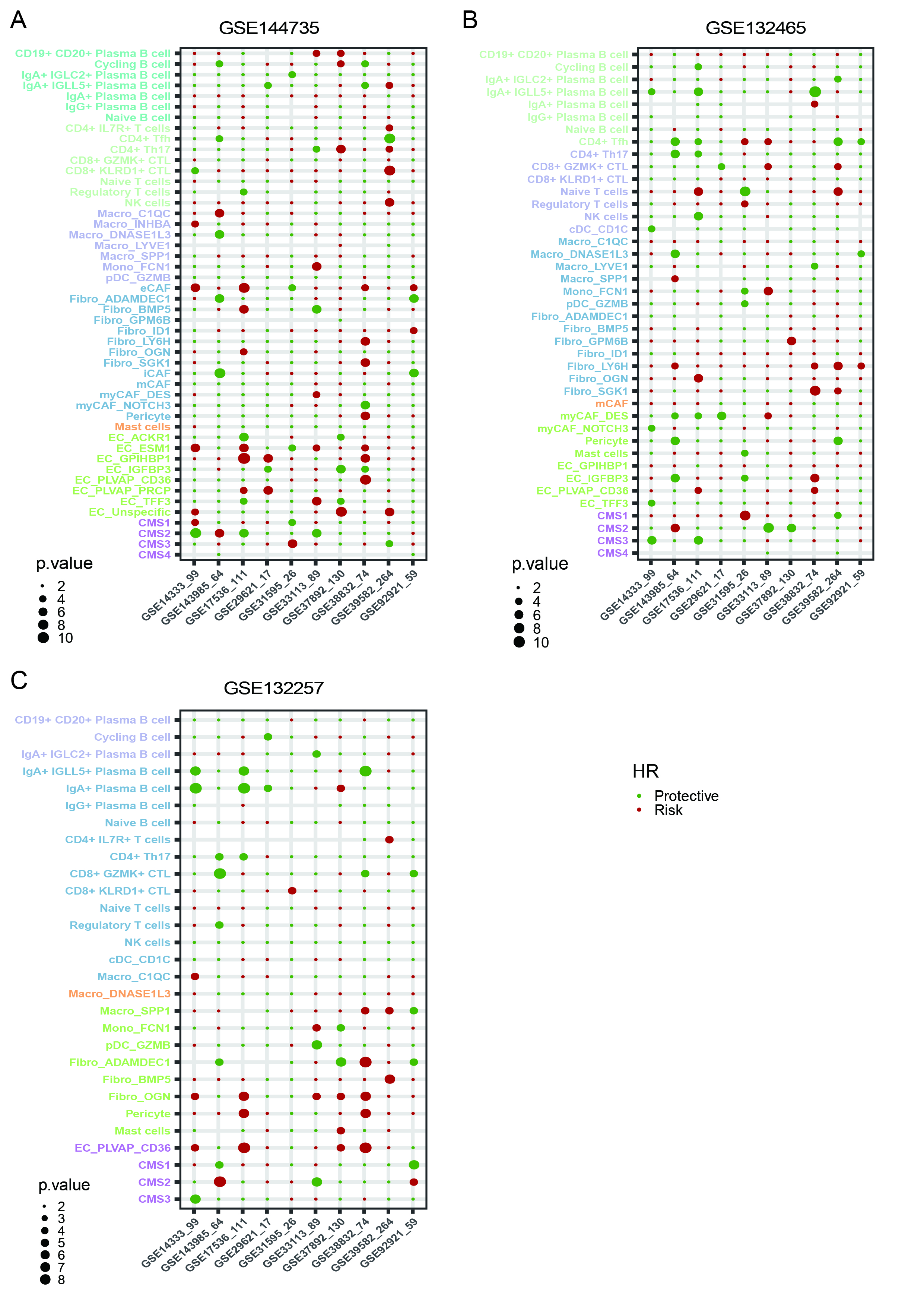

### Supplemental Figure13

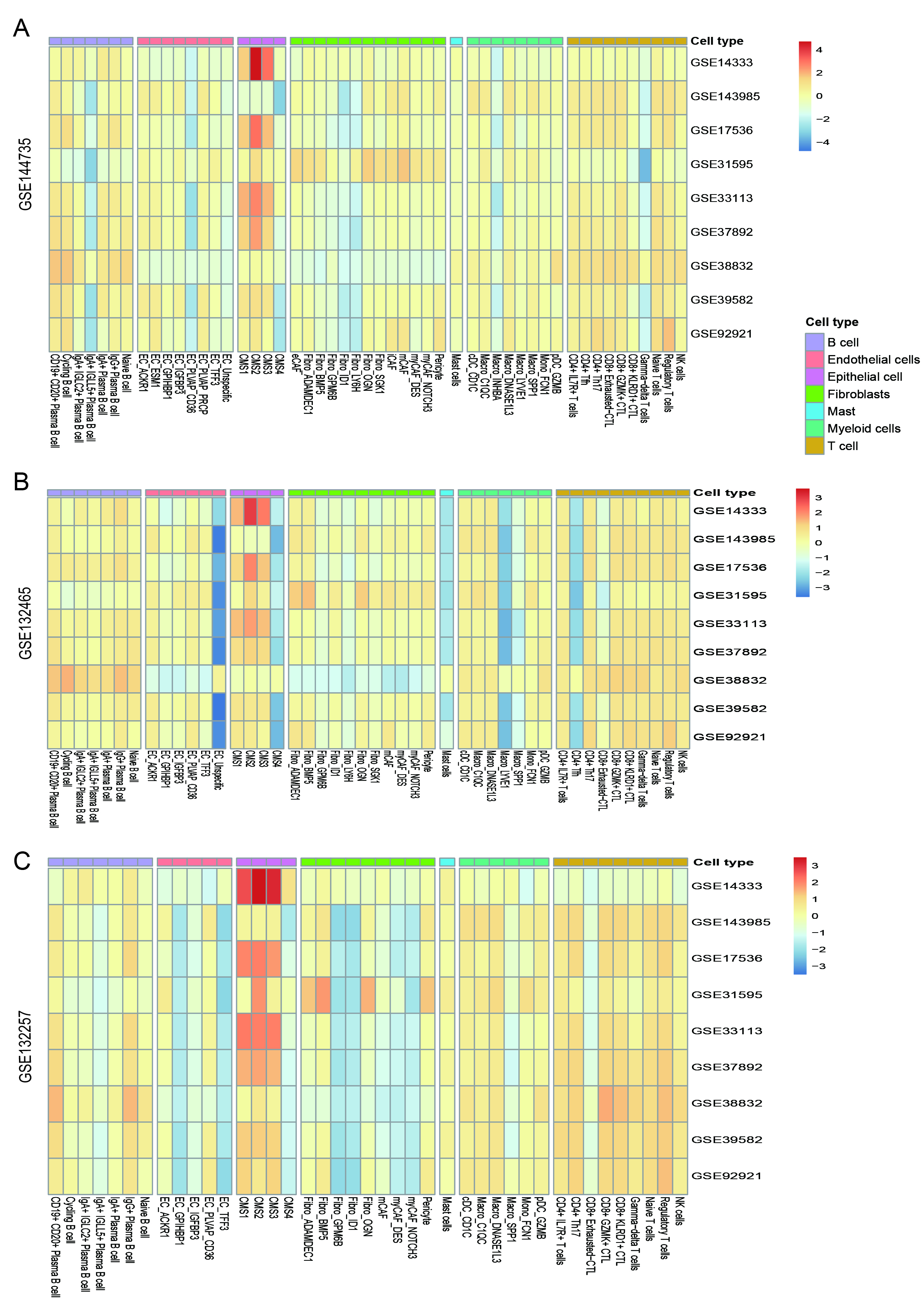

### Supplemental Figure14

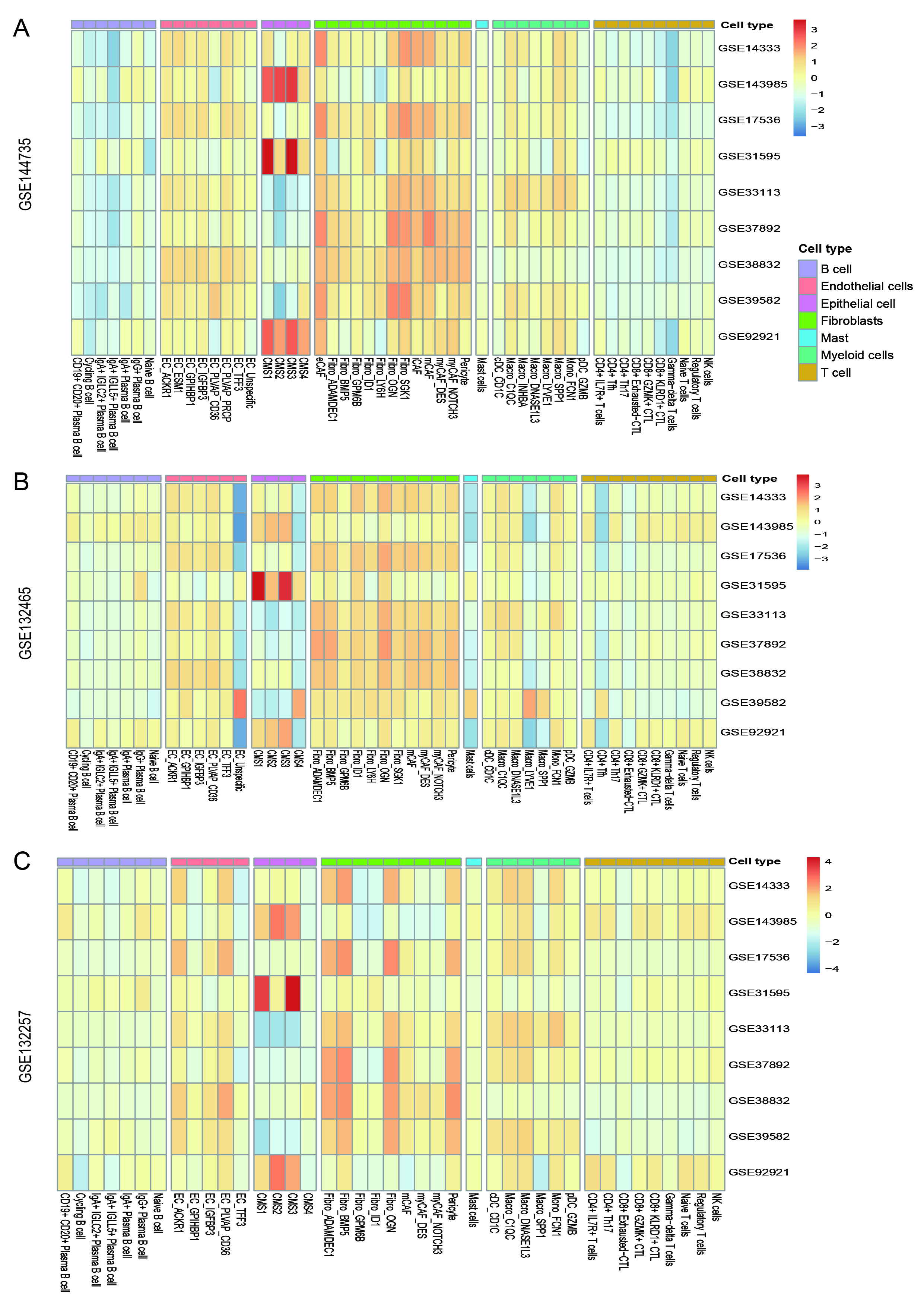

### Supplemental Figure15

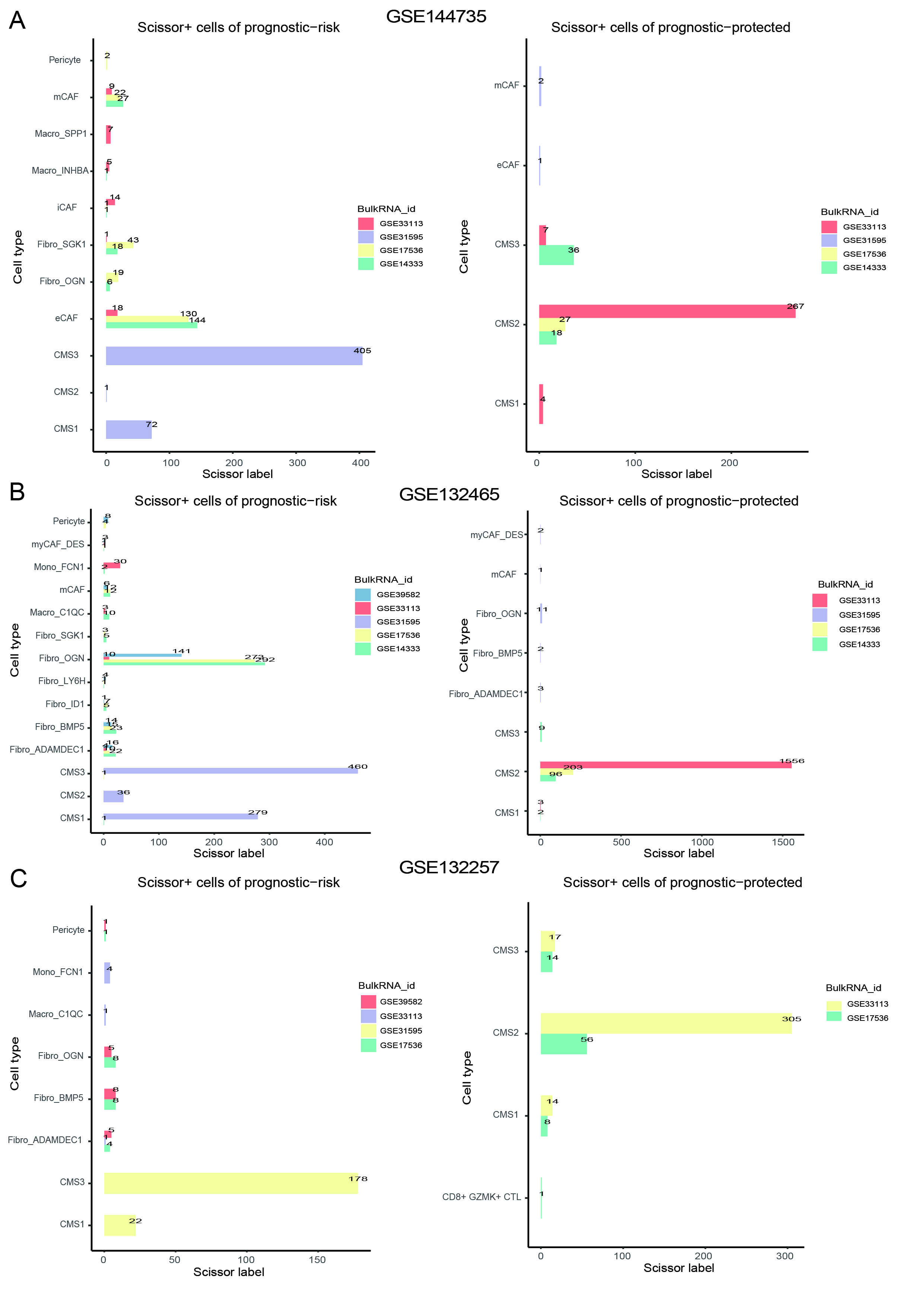

### Supplemental Figure16

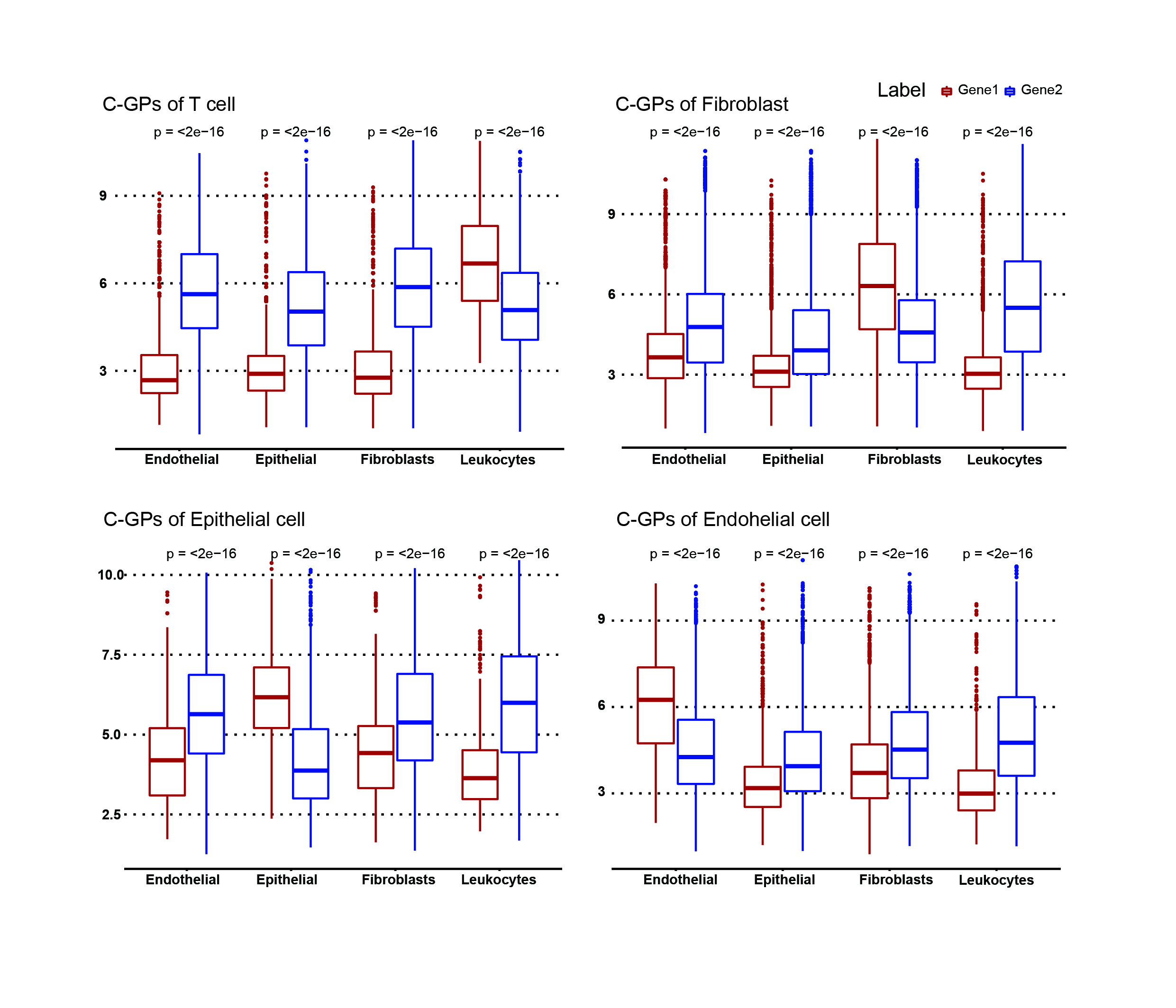

### Supplemental Figure17

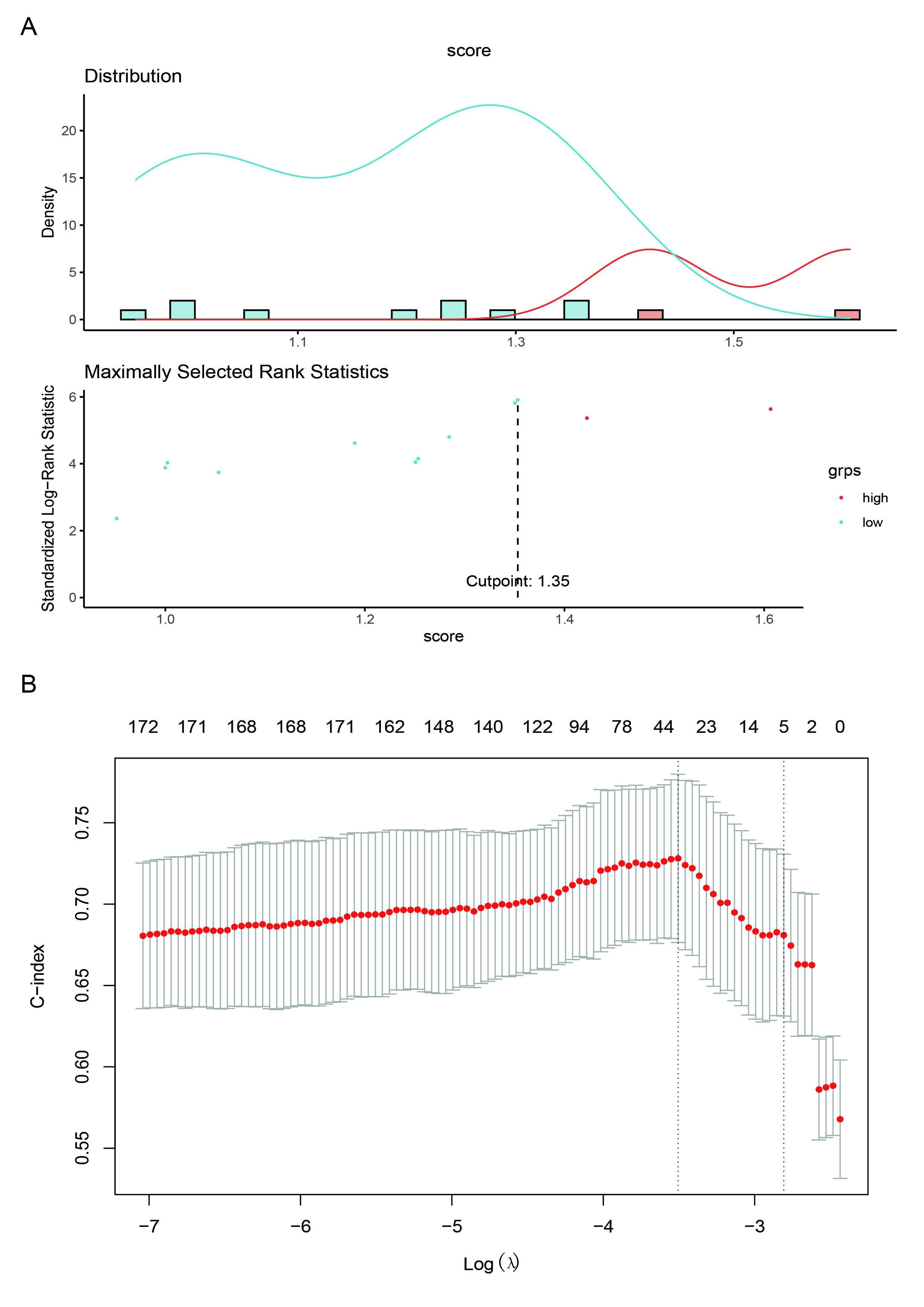

### Supplemental Figure18

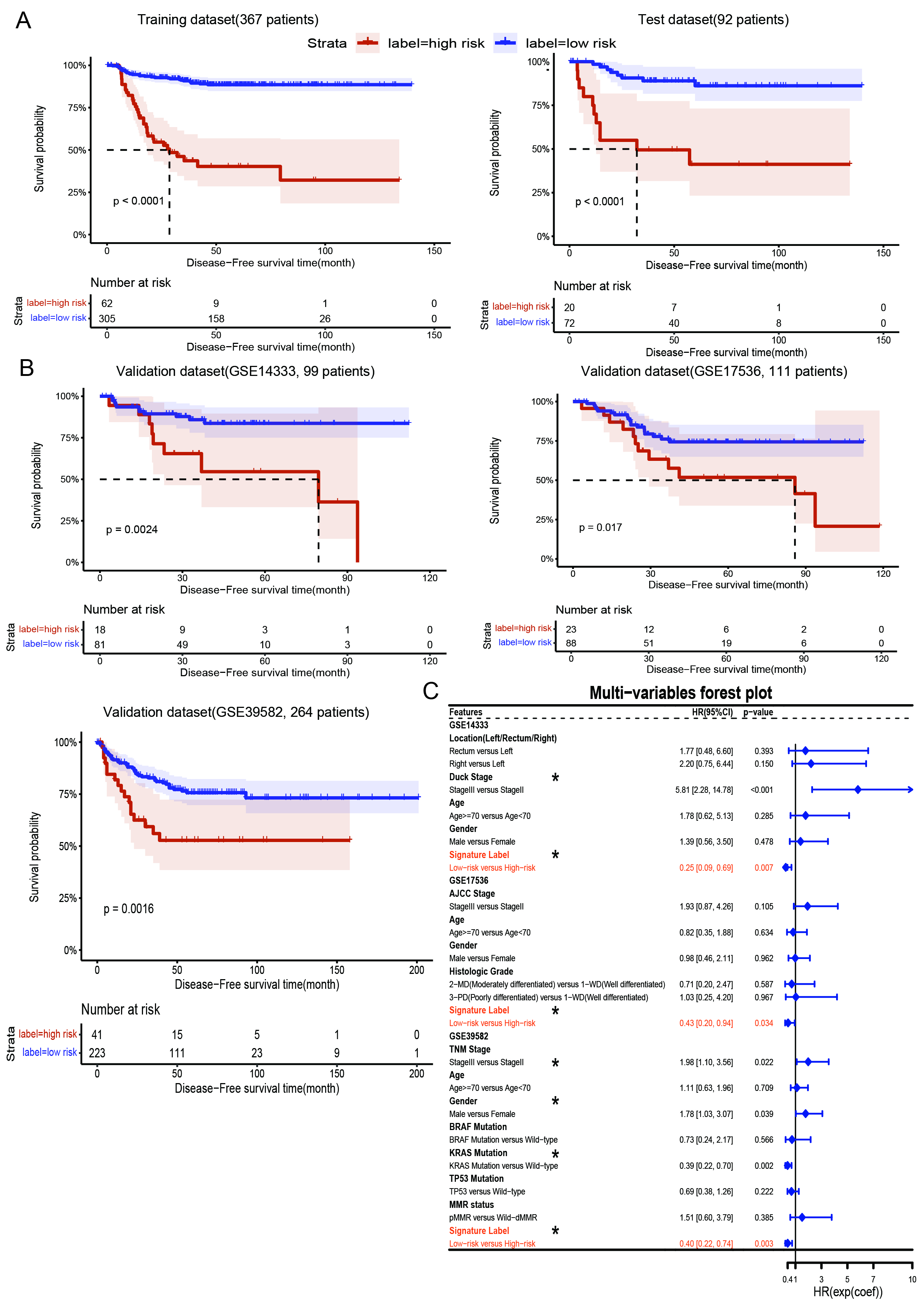

### Supplemental Figure19

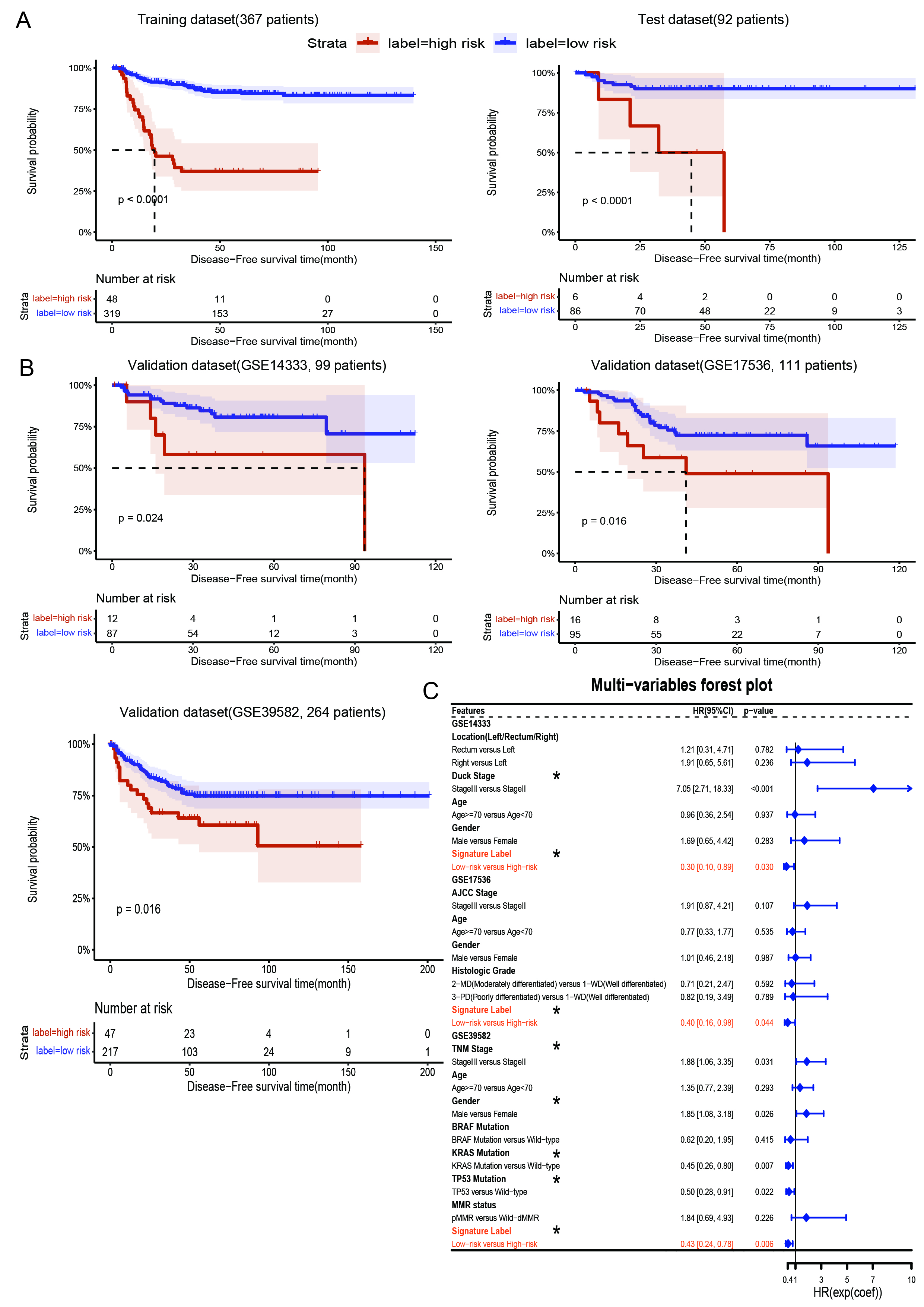
